# MiRNA-501-3p and MiRNA-502-3p: A Promising Biomarker Panel for Alzheimer’s Disease

**DOI:** 10.1101/2025.01.09.632227

**Authors:** Davin Devara, Bhupender Sharma, Gunjan Goyal, Daniela Rodarte, Aditi Kulkarni, Nathan Tinu, Ayana Pai, Subodh Kumar

## Abstract

**INTRODUCTION:** Alzheimer’s disease (AD) lacks a less invasive and early detectable biomarker. Here, we investigated the biomarker potential of miR-501-3p and miR-502-3p using different AD sources.

**METHODS:** MiR-501-3p and miR-502-3p expressions were evaluated in AD CSF exosomes, serum exosomes, familial and sporadic AD fibroblasts and B-lymphocytes by qRT-PCR analysis. Further, miR-501-3p and miR-502-3p expressions were analyzed in APP, Tau cells and media exosomes.

**RESULTS:** MiR-501-3p and miR-502-3p expressions were significantly upregulated in AD CSF exosomes relative to controls. MiRNA levels were high in accordance with amyloid plaque and NFT density in multiple brain regions. Similarly, both miRNAs were elevated in AD and MCI serum exosomes compared to controls. MiR-502-3p expression was high in fAD and sAD B-lymphocytes. Finally, miR-501-3p and miR-502-3p expression were elevated intracellularly and secreted extracellularly in response to APP and Tau pathology.

**DISCUSSION:** These results suggest that miR-501-3p and miR-502-3p could be promising biomarkers for AD.

## BACKGROUND

Alzheimer’s disease (AD) is a progressive, neurodegenerative disorder that remains the most common cause of dementia, accounting for 60-80% of all cases.^1^ Approximately 6.7 million people in the United States over the age of 65 are currently living with AD. By 2060, this is projected to grow to 13.8 million.^1^ In AD, amyloid beta (Aβ) plaques, composed of Aβ peptides, and neurofibrillary tangles (NFTs), composed of hyperphosphorylated tau (p-tau), accumulate in the brain and trigger proinflammatory events that cause neurodegeneration.^2,3^ The basal forebrain cholinergic neurons are among the affected cells, which causes an acetylcholine deficiency. This plays a major role in cognitive decline and is considered to be a hallmark of AD.^4,5^ Currently, no precise biomarkers or curative treatment exists for AD. FDA-approved medications remain limited, and most only provide temporary symptomatic benefits.^6–9^

One of the diagnostic tools we have for AD is the cerebrospinal fluid (CSF) analysis of p-tau, Aβ42, and total tau protein content, which reflects amyloidosis and tauopathy in the brain.^10^ However, for a successful diagnosis, the disease would already have to be relatively advanced.^2^ Early diagnosis of AD leads to proactive treatment of the disease, thus delaying the onset of dementia. This will not only improve prognosis, but it will also result in substantial cost savings to healthcare systems.^11^ For these reasons, searching for novel biomarkers has been a large focus of AD research in recent history.^12,13^ One of the most promising biomarkers that have emerged is microRNA (miRNA).^3,14–18^ The role of miRNAs as biomarkers is currently being studied in many diseases, including cancers, autoimmune conditions, and neurodegenerative disorders.^3,17,18,19–25^ Many miRNAs have been found to have differential expression in serum and CSF of AD patients, suggesting their possible potential as biomarkers.^26^ Among them, we had previously studied miR-455-3p extensively regarding its role in AD and its potential as a peripheral biomarker.^27–31^ Despite the numerous candidates, none of these potential miRNA biomarkers have yet passed the clinical trials stage. Therefore, it is important to continue developing our understanding of the importance of miRNAs in a clinical setting.

Recently, our lab discovered the high fold expression of microRNA-501-3p (miR-501-3p) and microRNA-502-3p (miR-502-3p) in the synaptosomes (synapses) of AD brains relative to unaffected controls.^32,33^ In this study, we found that miR-501-3p and miR-502-3p were among the top miRNAs sequentially overexpressed as AD progressed through the Braak stages. We also confirmed high expression levels of miR-501-3p and miR-502-3p in the synaptosomes of APP and Tau transgenic mice.^32^ Further characterization of these two synapse-specific miRNAs showed that they regulate the GABAergic synaptic pathway by reducing *GABRA1* expression. Disruption of this pathway plays a role in the synaptic dysregulation seen in AD, which was further supported by the increasing expression of miR-501-3p and miR-502-3p with worsening AD stages.^32^ Our recent findings investigated the roles of miR-502-3p in the modulation of GABAergic synapse function in AD.^34,35^ Following, we further explored the possible roles of miR-501-3p and miR-502-3p in various human diseases, including AD.^3^ Since these miRNAs are differentially expressed in many disorders, they could potentially serve as biomarkers for AD. Therefore, it is important to determine the status of miR-501-3p and miR-502-3p in the periphery, such as CSF or serum, to elucidate if differential levels of these miRNAs could reflect AD progression. Here, we aim to explore the biomarker potential of miR-501-3p and miR-502-3p by assessing their presence in central and peripheral circulation. We analyzed them in AD CSF, serum, fibroblasts, and B-lymphocytes. We also investigated the cellular and extracellular status of miR-501-3p and miR-502-3p in response to Aβ and tau pathology using cell culture studies. Our study on various AD sources unveiled strong biomarker capabilities of miR-501-3p and miR-502-3p for AD.

## METHODS

### Cerebrospinal fluid samples

CSF samples from AD patients and unaffected controls (UC) were obtained from NIH NeuroBioBank Center-Mount Sinai NIH Brain and Tissue Repository, 130 West Kingsbridge Road Bronx, NY59. Thirty-six CSF samples, including AD patients (n=26), age and sex-matched UC samples (n=10), were received from NIH NeuroBioBank. Demographic details such as: age, tissue type (CSF/ Serum), manner of death (Normal/ Suicide), sex, race, age, PMI, and diagnosis of the AD and UC CSF samples are summarized in **Supplementary Table 1.** The samples were collected during the period of 1988 to 2022. We received the neuropathology reports from the NIH NeuroBioBank that corresponded with each CSF sample used in our study. The neuropathological details of the samples such as plaques and tangle density in the different brain regions are summarized in **Supplementary Table 2**. The amyloid plaques and neurofibrillary tangle (NFT) density were determined in the entorhinal cortex, amygdala, hippocampus, middle frontal gyrus, superior temporal gyrus and inferior parietal lobule areas following neuropathology reports. The plaques and tangles distribution are categorized as none, sparse, moderate, frequent/severe based on the plaques and tangle density. The study was conducted at the Center of Emphasis in Neuroscience, Department of Molecular and Translational Medicine, Texas Tech University Health Sciences Center, El Paso, and Institutional Biosafety Committee (IBC protocol #22008) approved the study protocol for the use of human CSF specimens obtained from NIH NeuroBioBanks. The NIH NeuroBioBanks mentioned above are operated under their institution’s IRB approval, and they obtained written informed consent from the donors.

### Serum samples

Serum samples from AD patients and UC were obtained from Texas Alzheimer’s Research and Care Consortium (TARCC). Fifty-nine serum samples, including AD patients (n=25), mild cognitive impairment (n=15), and age and sex-matched UC samples (n=19), were received from TARCC. **Supplementary Table 3** summarizes the samples’ age, sex, race, Mini-Mental State Examination (MMSE) score, APOE information (when available), and total tau (when available).

### Fibroblasts and B-lymphocyte samples

Fibroblasts and B-lymphocyte samples were obtained from the Coriell Institute of Medical Research, 403 Haddon Ave, Camden, NJ. Three levels of disease pathology differentiation existed within the fibroblast and B-lymphocyte samples: familial AD (fAD), sporadic AD (sAD), and UC samples. Eighteen fibroblast samples, including fAD (n=4), sAD (n=6), and UC (n=8) were received from the Coriell Institute of Medical Research. Additionally, we received twenty-two B-lymphocyte samples, including fAD (n=6), sAD (n=6), and UC (n=10) were received from the from the same institution. Demographic information for fibroblast and B-lymphocyte samples, such as sex, age, biopsy source, tissue type, race, and disease status are summarized in **Supplementary Table 4** and **Supplementary Table 5,** respectively.

### SH-SY5Y cells and plasmid transfection

Human neuroblastoma (SH-SY5Y) cells are routinely maintained in our lab (Kumar et al., 2019). Cells were cultured in Dulbecco’s modified eagle medium (DMEM; Gibco^TM^) supplemented with Ham’s F-12 nutrient media, 10% fetal bovine serum (exosome-depleted FBS) and 1X antibiotic-antimycotic (Gibco) at 37^°^C in a humidified atmosphere containing 5% CO2. Cells were transfected with control plasmid (VB010000-9829sne; Vector builder), APP plasmid (pCAX APP Swe/Ind; AddgGene), and Tau plasmid (pRK5-EGFp-Tau E14 P301L; AddGene) using Lipofectamine 3000 following the manufacturer protocol (ThermoFisher Scientific, USA). The complete details of plasmids are provided in **Supplementary Information File 1**. At 48 hours post-transfection, the spent media (3 mL) were collected from control, APP and Tau plasmids transfected cells for exosome isolation.

### Exosome isolation

Exosomes were extracted from CSF, serum and cell culture media using specific kits following the manufacturer’s instructions. Exosomes were isolated from CSF samples using the total exosome isolation reagent (from other body fluids) kit (Catalog Number: 4484453; Invitrogen, USA). Serum exosomes were isolated from the serum samples using total exosome isolation reagent (from serum) kit (Catalog Number: 4478360) (Invitrogen, USA). Exosomes from the cell culture media were isolated by using the total exosome isolation reagent (from cell culture media) kit (Catalog Number: 4478359; Invitrogen, USA).

### Transmission electron microscopy of exosomes

Exosome were isolated from CSF, serum, cell culture media and their morphology were characterized by transmission electron microscopy (TEM) analysis (Kumar et al., 2021). Exosome pellet was washed with 1XPBS and dissolved in a fixative solution (8% Glutaraldehyde, 16% paraformaldehyde, and 0.2 M sodium cacodylate buffer) for 1 h at room temperature. Samples were centrifuged at 300g for 10 minutes and the resulting exosome pellet underwent electron microscopy at the imaging core facility at Texas Tech University, Lubbock, Texas, US.^31^

### Immunoblotting analysis for exosome marker proteins

For immunoblotting analysis, exosomes pellets isolated from CSF, serum and cell media were suspended in RIPA buffer (Thermo scientific) with (1X) protease inhibitor and EDTA. The samples were sonicated (Amplitude 50%, Pulse 2 sec on/off) for ten seconds on ice. Exosome proteins were quantified by bicinchoninic acid assay (BCA) method. The 40 μg of exosome protein were mixed in a 4:1 (v/ v) ratio with 4x LDS sample loading buffer (Novex) and subjected to SDS-PAGE analysis. The electrophoresis was carried out in mini protein tetra cell (BioRad) at 100 V for 1 hour. Transfer sandwich was prepared with an anode-facing gel and a cathode-facing PVDF membrane. Protein transfer was carried out for 10 minutes in transfer buffer using the trans-blot turbo (BioRad) transfer system. Thereafter blot (PVDF-membrane) was blocked with 5% (w/ v) bovine serum albumin (BSA) in tris-buffer saline with 0.1% tween (1X TBS-T). Blot(s) were incubated overnight with primary antibodies specific to anti-human biomarkers *i.e.* CD9 (Mouse; 1:500), CD63 (Mouse; 1:500), TSG101 (Mouse; 1:500), NeuN (Mouse; 1:2000), GFAP (Mouse; 1:2000), IBA1 (Mouse; 1:2000), APP (Rabbit; 1:1000), Tau (Mouse; 1:1000) and GAPDH (Rabbit; 1:3000) prepared in BSA (5% w/ v; 10 mL) at 4^°^C. The complete details of antibodies and their dilutions are provided in **Supplementary Table 6**. Thereafter blot(s) were washed three times with TBS-T and incubated for two hours at room temperature with secondary antibodies *i.e.* anti-mouse/ anti-rabbit IgG-peroxidase (Sigma; 1:10,000 prepared in BSA 5% w/ v). After washing the blot(s) five times with TBS-T, the protein bands labeled with secondary antibodies were detected using the Clarity™ enhanced chemiluminescence (ECL; Bio-Rad, USA) western blotting substrate. Blot(s) were then visualized by exposing it to dark conditions in luminescent image analyzer (Amersham imager 680; GE Healthcare Bio-Sciences, Sweden).^31^

### Particle size analysis of exosomes

The particle size of micro-vesicles isolated from CSF, serum, and cell culture media were determined by Dynamic Light Scattering (DLS) analysis using the NanoSight (Malvern zetasizer, Worcerestershire, UK). The exosomes pellet was diluted with 1X phosphate buffer saline (1:100) and processed for exosomes size distribution analysis. The particle size and density were recorded along all the sample groups.^36^

### Exosome RNA isolation

The exosome pellets from CSF, serum, cell media were suspended in Trizol (500 μL) for total RNA isolation. Chloroform (100 μL) was added to each sample, vortexed and incubated at room temperature for 10 minutes. Samples were centrifuged at 12,000 g for 30 minutes at 4^°^C. Aqueous phase was separated out from each sample(s) into fresh eppendorf(s). Thereafter, isopropanol (250 μL) was added to each sample(s) and incubated overnight at –20^°^C. RNA pellets were collected by centrifugation at 12,000 g for 30 minutes at 4^°^C. Thereafter pellet(s) were washed with ethanol (500 μL; 75% v/v). Samples were again centrifuged at 12,000 g for 30 minutes at 4^°^C. After washing ethanol was removed and samples were air dried and suspended in DEPC-treated water (12 μL). The concentration (ng/ µL) of RNA was determined by nano-drop 2000 spectrophotometer (Thermo scientific; USA). The cDNA was synthesized from total RNA including miRNA in two steps firstly the polyadenylation of the miRNA followed by the cDNA synthesis using 1^st^ strand cDNA synthesis kit (Agilent technologies, USA; Catalog Number: 600036) in the thermal cycler (Applied biosystem 9902; US) according to the manufacturer’s protocol.^31^

### Quantitative real-time PCR analysis of miR-501-3p and miR-502-3p

The expression of miR-502-3p and miR-501-3p were quantified by qRT-PCR. qRT-PCR was performed by preparing a reaction mixture containing 1 μL of miRNA-specific forward primer (10 μm), 1 μL of a universal reverse primer (3.125 μM) (Agilent Technologies Inc., CA, USA), 5 μL of 2XSYBR green PCR master mix (Kappa SYBR), and 1 μL of cDNA. To this mixture, RNase-free water was added to a total of 10 μL final volume. Primers used in the current study were synthesized commercially (Integrated DNA Technologies, Inc., City, Iowa, USA) for miR-502-3p, miR-501-3p and U6 small nuclear RNA **(Supplementary Table 7)**. To normalize the miRNA expression U6 snRNA expression was used as an internal control. The reaction mixture for each sample was prepared in duplicates and set in the Roche Real-time PCR system (LightCycler instrument 96 Instrument; US). MiRNA fold changes were calculated by using the formula (2^− ΔΔct^).^28,29,31,37^

### MiR-501-3p and MiR-502-3p analysis with amyloid plaques and NFT density in affected brain regions

We evaluated the neuropathology report of AD CSF samples in relation to amyloid plaques and NFT density across affected brain regions. The amyloid plaques and NFT densities were categorized as none, sparse, moderate, and frequent/severe. The percentage of sample distribution and miR-501-3p and miR-502-3p fold changes were accessed in the entorhinal cortex, hippocampus, amygdala, middle frontal gyrus, inferior parietal lobule, and superior temporal gyrus brain region. Further, miR-502-3p and miR-501-3p fold variations were determined in the samples with sparse, moderate, and frequent/severe amyloid plaques and NFT compared to none.

### Statistical analysis

Statistical parameters were calculated using GraphPad Prism software, v8 (La Jolla, CA, USA) and OriginPro 2024b (Northampton, MA, USA). Results are reported as mean ± SD and median ± SEM. The QRT-PCR results were analyzed by students t tests (unpaired t test) comparing the miR-501-3p and miR-502-3p expression in AD versus (vs.) UC CSF exosomes. The ordinary one-way ANOVA was implied to compare the miR-501-3p and miR-502-3p expression in UC serum vs. MCI serum vs. AD serum exosomes. The unpaired t test was applied to compare the miR-501-3p and miR-502-3p expression in UC fibroblast vs. fAD fibroblasts and sAD fibroblasts. Similarly, unpaired t test was applied to compare the miR-501-3p and miR-502-3p expression in UC B-lymphocytes vs. fAD B-lymphocytes and sAD B-lymphocytes. Further, unpaired t test was used to compare the miR-501-3p and miR-502-3p expression in control vs. APP and Control vs. Tau cells. P < 0.05 was considered statistically significant.

## RESULTS

We have divided the present investigation into five different studies. **Figure 1A** briefly outlined the overall work plan of the study. We used five different AD sources to investigate the biomarker potential of miR-501-3p and miR-502-3p in relation to amyloid beta and tau pathology: (1) CSF, (2) serum, (3) fibroblasts, (4) B-lymphocytes, and (5) APP and Tau plasmid transfected cells.

**Figure 1.**
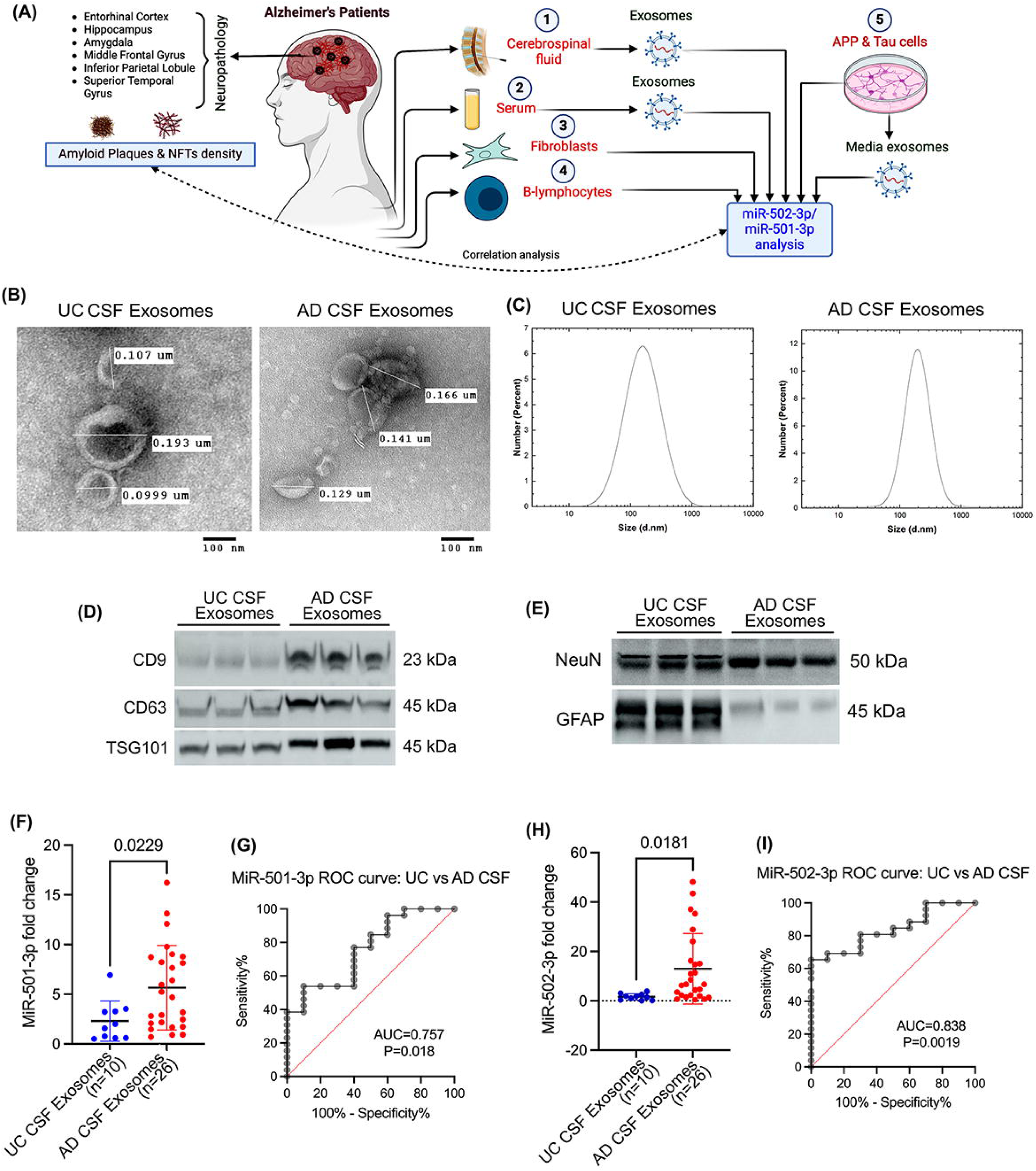
Status of miR-501-3p and miR-502-3p in AD and UC CSF samples. **(A)** Brief study plan: Exosomes were extracted from AD and UC CSF, serum, APP and tau transfected cells. Afterward, miR-501-3p and miR-502-3p expression were quantified from these exosomes. Additionally, miR-501-3p and miR-502-3p quantification was also performed from AD and UC fibroblasts and B-lymphocytes. Expression of these two miRNAs from CSF samples was then correlated with the neuropathology report of the same samples regarding the amyloid plaques and NFT density in different brain regions. **(B)** TEM images of exosomes from AD and UC CSF samples at 100 nm magnification. **(C)** Particle size analysis of exosomes using DLS. **(D)** Immunoblotting of exosome marker proteins CD9, CD63, and TSG101. **(E)** Immunoblotting of neuronal marker, NeuN, and astrocytic marker, GFAP. **(F)** Expression of miR-501-3p in UC CSF exosomes (n=10) compared with AD CSF exosomes (n=26) (t-test, p=0.0229). **(G)** ROC curve analysis of miR-501-3p expression in AD relative to UC CSF exosomes (AUC=0.757, p=0.018). **(H)** Expression of miR-502-3p in UC CSF exosomes (n=10) compared with AD CSF exosomes (n=26) (t-test, p=0.0181). **(I)** ROC curve analysis of miR-502-3p expression in AD relative to UC CSF exosomes (AUC=0.838, p=0.0019).

### 1. Cerebrospinal Fluid Study

#### Characterization of CSF Exosomes in UC and AD Samples

We begin by evaluating CSF samples obtained from UC and AD patients. First, the exosomes were isolated from the CSF samples. The isolated exosomes were then characterized by TEM, immunoblotting, and DLS analysis. The TEM analysis showed the cup-shaped ultrastructure of exosomes in both UC and AD CSF samples **(Figure 1B)**. The exosome vesicle size ranged from

0.101 micrometers (μm) to 0.185 (μm) in both UC and AD. We did not see any significant size differences between the exosomes in UC vs. AD. Next, we determined the micro-vesicle size range using the DLS analysis. The UC CSF exosomes ranged from 10-110 nm with peaks of around 40 nm, and AD CSF exosomes ranged from 10-300 nm with peaks of around 80 nm **(Figure 1C)**. Next, we characterized the exosomes by assessing the expression of exosome marker proteins CD9, CD63, and TSG101. The immunoblotting analysis showed detectable levels of CD9, CD63, and TSG101 proteins **(Figure 1D)**. Furthermore, we aimed to determine which brain cells produced the isolated exosomes. Immunoblotting analysis of brain cell markers showed a detectable level of neuronal marker NeuN and astrocyte marker GFAP **(Figure 1E)**. We performed the immunoblotting analysis of microglial marker IBA1 but did not see detectable expression of IBA in CSF exosomes. These results confirmed successful exosome extraction from the CSF and suggested that the exosomes released into the CSF contributed by both neurons and astrocytes.

#### MiR-501-3p and MiR-502-3p Expression Analysis in CSF Exosome

To determine the biomarker potential of miR-501-3p and miR-502-3p in CSF, we assessed their expression levels in the exosomes isolated from UC and AD CSF samples. QRT-PCR analysis of miR-501-3p showed a significantly (p=0.0229) higher expression of miR-501-3p in AD CSF exosomes relative to UC CSF exosomes **(Figure 1F)**. Afterward, to assess the diagnostic value of miR-501-3p fold-change in AD vs. UC samples, we conducted a receiver operating characteristic (ROC) curve analysis. Our results showed a significant area under the curve (AUC) value for miR-501-3p fold change (AUC=0.757, p=0.018) **(Figure 1G)**.

Similarly, we analyzed the miR-502-3p expression in UC and AD CSF exosome samples. QRT-PCR analysis of miR-502-3p also showed a significantly (p= 0.0181) higher expression of miR-502-3p in AD CSF exosomes relative to UC CSF exosomes **(Figure 1H)**. We conducted a ROC curve analysis for miR-502-3p fold change, which indicated a significant AUC value for miR-502-3p fold change (AUC=0.838; P=0.0019) in AD CSF exosomes vs. UC CSF exosomes **(Figure 1I)**. These observations suggest that both miRNAs could be a potential synapse-associated exosome biomarker for AD and showed its discriminating power between AD and UC.

#### MiR-501-3p Expression Relative to Amyloid Plaque and NFT Density in Affected Brain Regions

We wanted to investigate if there was a relationship between CSF miRNA fold change and AD pathology. To do so, we first compared miR-501-3p fold change with neuropathology report of amyloid plaque density in the entorhinal cortex, amygdala, hippocampus, middle frontal gyrus, and inferior parietal lobule of the same patients **(Figure 2A-E).** Since there was a vastly variable distribution of patients in the different severity categories, we could only make qualitative observations. A majority (69%) of the patients had moderate (n=7) or severe (n=11) amyloid plaque density in the entorhinal cortex **(Figure 2A).** Aside from the sparse severity, the data shows a positive correlation between average miR-501-3p fold change and amyloid plaque density in the entorhinal cortex **(Figure 2A).** Similarly, 85% of patients had moderate (n=11) or severe (n=10) amyloid plaque density in the amygdala **(Figure 2B).** No correlation can be observed between miR-501-3p fold change and plaque severity in the amygdala. Interestingly, only 46% of patients showed moderate (n=9) or severe (n=3) amyloid plaque density in the hippocampus **(Figure 2C).** Here, miR-501-3p fold change is most highly correlated with sparse amyloid plaque density **(Figure 2C).** Next, 54% of patients had severe amyloid plaque density (n=19), and 34% of patients had no amyloid plaque density (n=12) in the middle frontal gyrus **(Figure 2D).** Here, miR-501-3p fold change is most highly correlated with severe amyloid plaque density **(Figure 2D).** Finally, 62% of patients had moderate (n=7) or severe (n=15) amyloid plaque density in the inferior parietal lobule, while 34% had no plaques (n=12) **(Figure 2E).** For this brain region, miR-501-3p fold change is most highly correlated with severe amyloid plaque density **(Figure 2E).**

**Figure 2.**
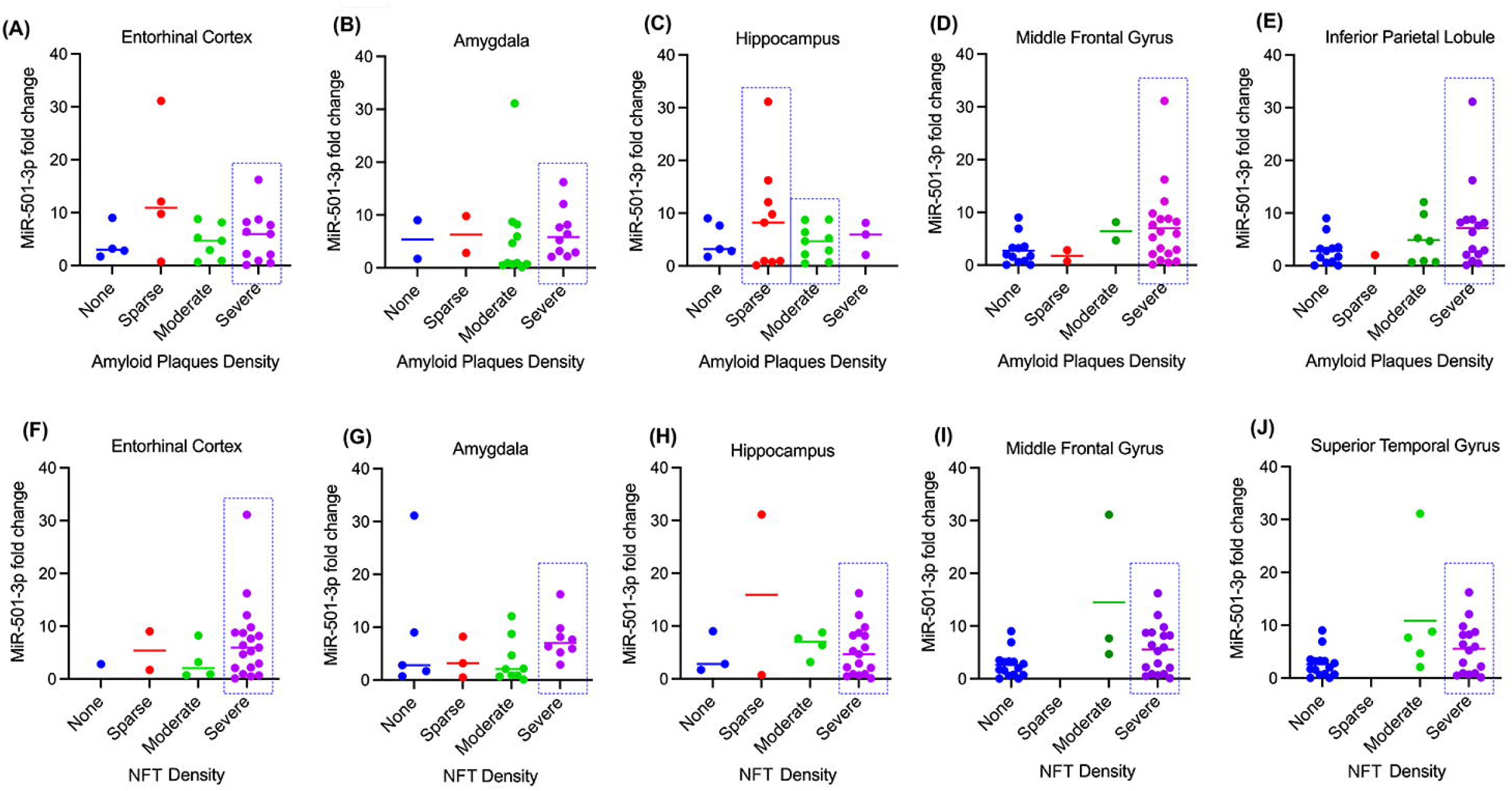
Relationship of miR-501-3p expression and the neuropathology in AD CSF samples. MiR-501-3p expression was correlated with amyloid plaque density in the **(A)** entorhinal cortex, **(B)** amygdala, **(C)** hippocampus, **(D)** middle frontal gyrus, and **(E)** inferior parietal lobule. MiR-501-3p expression was also correlated with NFT density in the **(F)** entorhinal cortex, **(G)** amygdala, **(H)** hippocampus, **(I)** middle frontal gyrus, and **(J)** superior temporal gyrus.

Next, we evaluated miR-501-3p level with NFT density in the entorhinal cortex, amygdala, hippocampus, middle frontal gyrus, and superior temporal gyrus in the AD brain **(Figure 2F-J).** 73% of patients had severe (n=19) NFT density in the entorhinal cortex **(Figure 2F).** While the data shows that miR-501-3p fold change is most highly correlated with severe NFT density, it is difficult to confirm this as there are very few patients in the other severity categories **(Figure 2F).** Next, a majority (68%) of the patients had moderate (n=9) or severe (n=8) NFT density in the amygdala **(Figure 2G).** Here, miR-501-3p fold change is most highly correlated with severe NFT density **(Figure 2G).** Unlike with amyloid plaque density, a majority (65%) of the patients had severe NFT density in the hippocampus **(Figure 2H).** It is difficult to conclude which severity has the highest correlation with miR-501-3p fold change here because there are few patients in the other categories. Similar to amyloid plaque density, 50% of patients had severe (n=18) NFT density in the middle frontal gyrus, and 42% of patients had no NFT density (n=15) **(Figure 2I).** This bimodal distribution of patients makes it difficult to conclude correlation patterns with miR-501-3p fold change and NFT density in this region of the brain. Finally, there was also a bimodal distribution of NFT density in the superior temporal gyrus, with 44% of patients having severe (n=16) NFT density and 42% of patients having no (n=15) NFT density **(Figure 2J).** As with the middle frontal gyrus, it is difficult to conclude correlation patterns with this distribution. Overall, these results suggest that miR-501-3p expression in the CSF could reflect some degree of amyloid plaque density or NFT density in the brain; however, a study with a higher number of samples that is more evenly distributed among the different severities is necessary to explore this further.

#### MiR-502-3p Expression Relative to Amyloid Plaque and NFT Density in Affected Brain Regions

Likewise, we also compared miR-502-3p fold change with amyloid plaque density in the entorhinal cortex, hippocampus, amygdala, middle frontal gyrus, and inferior parietal lobule of the same patients **(Figure 3A-E).** The distribution of amyloid plaque severity among the patients is the same as discussed in the previous section **(Figures 2A-E).** In the entorhinal cortex, miR-502-3p fold change average is better correlated with moderate than severe amyloid plaque density, but the patients that have the highest miR-502-3p fold change have severe amyloid plaque density **(Figure 3A).** In the amygdala, miR-502-3p fold change average is better correlated with severe amyloid plaque density than moderate **(Figure 3B).** Interestingly, miR-502-3p fold change is better correlated with sparse amyloid plaque density than moderate **(Figure 3C).** While the fold change for severe amyloid plaque density is higher on average than sparse, there are only 3 patients in that group. In the middle frontal gyrus, miR-502-3p fold change average is more correlated with severe amyloid plaque density than the other groups **(Figure 3D).** Finally, miR-502-3p fold change is more correlated with severe amyloid plaque density than the other groups in the inferior parietal lobule **(Figure 3E).**

**Figure 3.**
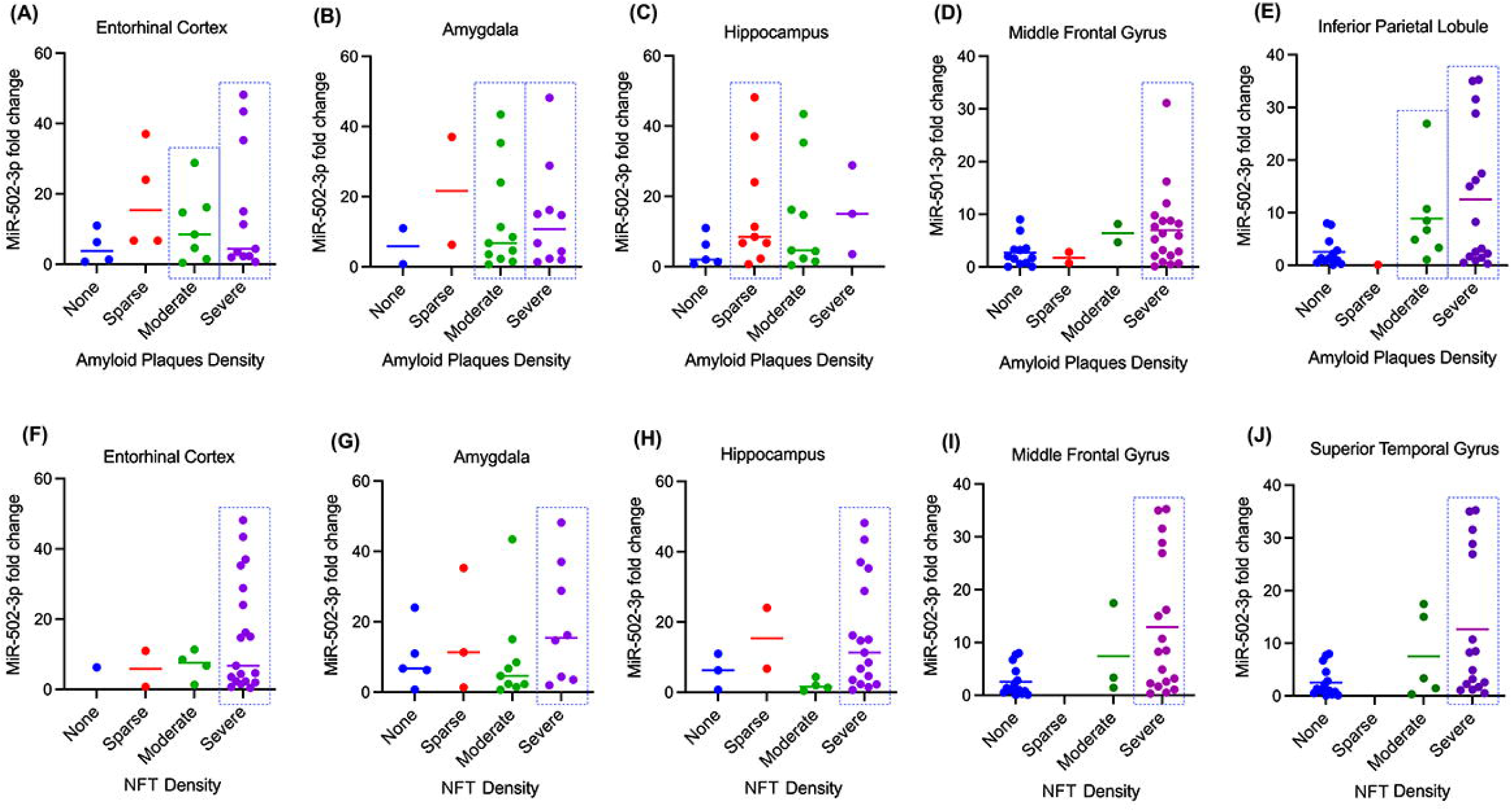
Relationship of miR-502-3p expression and the neuropathology in AD CSF samples. MiR-502-3p expression was correlated with amyloid plaque density in the **(A)** entorhinal cortex, **(B)** amygdala, **(C)** hippocampus, **(D)** middle frontal gyrus, and **(E)** inferior parietal lobule. MiR-502-3p expression was also correlated with NFT density in the **(F)** entorhinal cortex, **(G)** amygdala, **(H)** hippocampus, **(I)** middle frontal gyrus, and **(J)** superior temporal gyrus.

Next, we evaluated miR-502-3p level with NFT density in the entorhinal cortex, hippocampus, amygdala, middle frontal gyrus, and superior temporal gyrus in the AD brain **(Figure 3F-J).** Like for amyloid plaque density, the distribution of NFT density severities among the patients is the same as discussed in the previous section **(Figures 2F-J).** In the entorhinal cortex, the average miR-502-3p fold change is similar between the different NFT density severities; however, none, sparse, and moderate NFT density groups each have limited patients. As such, it is difficult to conclude the correlation here **(Figure 3F)**. In the amygdala, miR-502-3p fold change is more correlated with severe NFT density than the other groups **(Figure 3G).** In the hippocampus, miR-502-3p average fold change is better correlated with severe NFT density than moderate or no NFT density **(Figure 3H).** While miR-502-3p average fold change is higher in sparse NFT density than severe, there are only two patients in that group. Similarly, miR-502-3p average is most correlated with severe NFT density in both the middle frontal gyrus and superior temporal gyrus **(Figure 3I-J).**

Except for the entorhinal cortex, each brain region analyzed had severe amyloid plaque and NFT densities that most correlated with miR-502-3p average fold change. These results suggest that miR-502-3p expression in the CSF could reflect severe degrees of amyloid plaque density or NFT density in these brain regions. However, as stated in the section above, a study with a larger sample size must be conducted.

### 2. Serum Study

#### Characterization of Serum Exosomes in UC, MCI, and AD Samples

As with the CSF samples, the exosomes were isolated from AD, MCI, and UC serum samples. The isolated exosomes were characterized by TEM, immunoblotting and DLS analysis. The TEM analysis revealed the cup-shaped ultrastructure of exosomes in AD, MCI, and UC **(Figure 4A)**. Furthermore, the exosome vesicle size ranged from 0.110 μm to 0.119 μm in UC, 0.121 μm to 0.168 μm in MCI, and 0.210 μm to 0.270 μm in AD. Next, we tested for the expression of exosome marker proteins CD9, CD63, and TGS101. The immunoblotting analysis showed detectable CD9, CD63, and TGS101 protein levels in the UC, MCI, and AD samples **(Figure 4B)**. Next, we performed a DLS analysis to characterize the particle size further. The UC serum exosomes ranged from 10-110 nm with peaks of around 40 nm; MCI serum exosomes ranged from 10-200 nm with peaks of around 80 nm; and AD serum exosomes ranged from 10-300 nm with peaks of around 80 nm **(Figure 4C)**. Interestingly, on average, MCI serum exosomes and AD serum exosomes were larger than UC exosomes. Finally, to determine the origin of exosomes, we performed an immunoblotting analysis of several brain cell markers. This showed detectable levels of NeuN, GFAP, and IBA1, suggesting that these exosomes could have been released into the serum from neurons, astrocytes, and microglia **(Figure 4D)**. Overall, these results confirm the successful extraction of exosomes from the serum samples.

**Figure 4.**
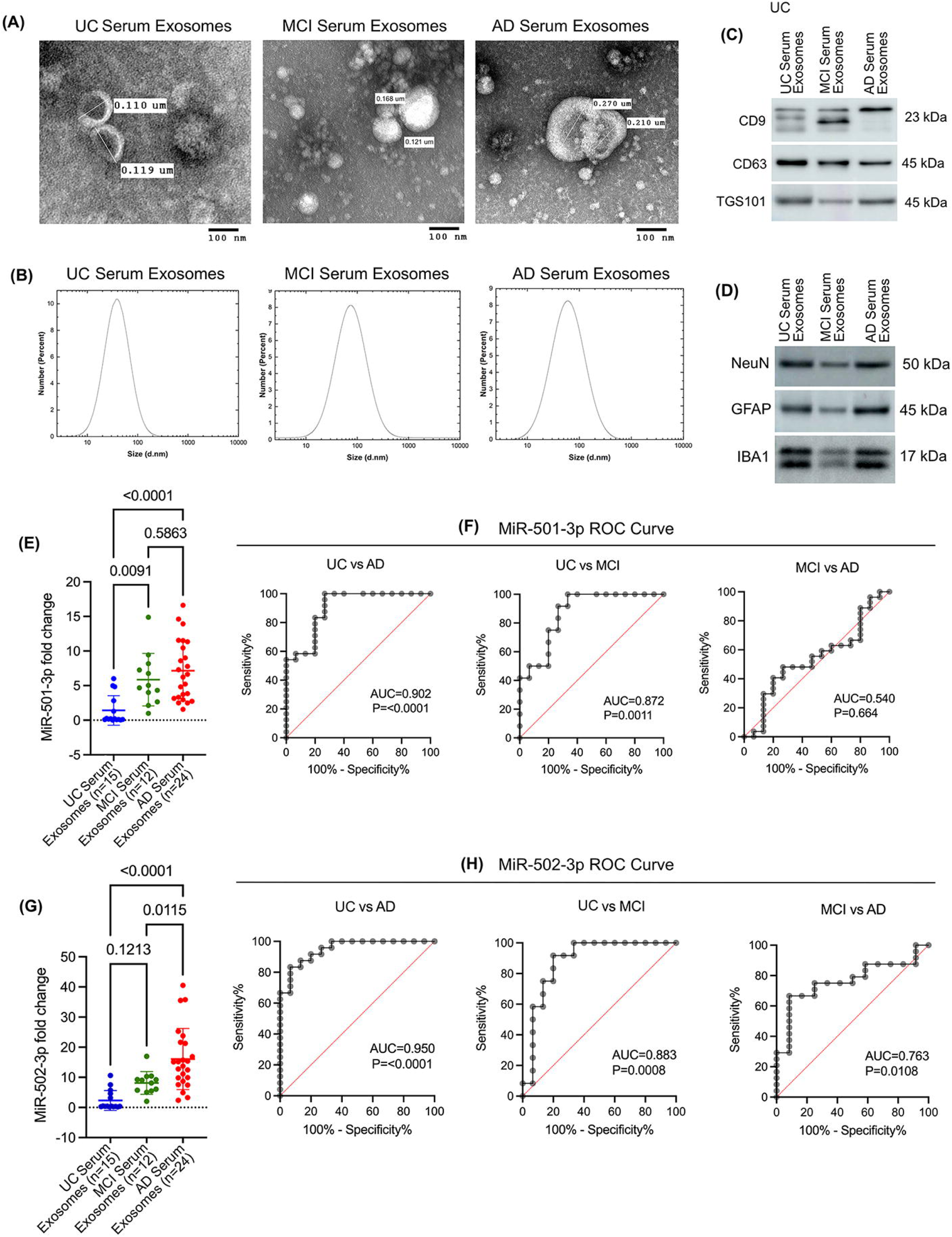
Characterization of miR-501-3p and miR-502-3p in UC, MCI, and AD serum samples. **(A)** TEM images of exosomes from UC, MCI, and AD serum samples at 100 nm magnification. **(B)** Particle size analysis of exosomes using DLS. **(C)** Immunoblotting of exosome marker proteins CD9, CD63, and TSG101. **(D)** Immunoblotting of neuronal marker, NeuN, and astrocytic marker, GFAP, and microglial marker, IBA1. **(E)** Expression of miR-501-3p in UC serum exosomes (n=15) compared with MCI serum exosomes (n=12, t-test, p=0.0091) and AD serum exosomes (n=24, one-way ANOVA, p<0.0001). Expression between MCI and AD serum exosomes was also compared (t-test, p=0.5863). **(F)** ROC curve analysis of miR-501-3p expression in UC vs. AD serum exosomes (AUC=0.902, p<0.0001), UC vs. MCI serum exosomes (AUC=0.872, p=0.0011), and MCI vs. AD serum exosomes (AUC=0.540, p=0.664). **(G)** Expression of miR-502-3p in UC serum exosomes (n=15) compared with MCI serum exosomes (n=12, t-test, p=0.1213) and AD serum exosomes (n=24, one way ANOVA, p<0.0001). Expression between MCI and AD serum exosomes was also compared (t-test, p=0.0115). **(H)** ROC curve analysis of miR-502-3p expression in UC vs. AD serum exosomes (AUC=0.950, p<0.0001), UC vs MCI serum exosomes (AUC=0.883, p=0.0008), and MCI vs. AD serum exosomes (AUC=0.763, p=0.0108).

#### MiR-501-3p and miR-502-3p expression Analysis in AD and MCI Serum Exosomes

To assess the potential of miR-501-3p and miR-502-3p as serum biomarkers, we evaluated their expression levels in the exosomes isolated from UC, MCI, and AD serum samples. QRT-PCR analysis showed higher levels of miR-502-3p and miR-501-3p in MCI serum exosomes relative to UC serum exosomes. Furthermore, we observed even higher levels of miR-502-3p and miR-501-3p in AD serum exosomes relative to MCI and UC serum exosomes. MiR-501-3p levels were significantly upregulated in MCI relative to UC (p=0.0091) and in AD relative to UC (p=<0.0001); however, miR-501-3p levels were not significantly higher in AD compared to MCI (p=0.5863) **(Figure 4E)**. Furthermore, we conducted a ROC curve analysis to assess the diagnostic value of miR-501-3p fold change in serum samples. Our results show a significant AUC for miR-501-3p fold change in MCI vs. UC (AUC=0.872; p=0.0011) and AD vs. UC (AUC=0.902; p=<0.0001) but not for AD vs. MCI (AUC=0.540; p=0.664) **(Figure 4F)**.

Interestingly, miR-502-3p levels were not significantly upregulated in MCI relative to UC (p=0.1213); however, they were significantly upregulated in AD relative to UC (p=<0.0001) and in AD compared to MCI (p=0.0115) **(Figure 4G)**. ROC analysis for miR-502-3p fold change showed a significant AUC for miR-502-3p fold change in MCI vs. UC (AUC=0.883; p=0.0008), AD vs. UC (AUC=0.950; p=<0.0001), and for AD vs. MCI (AUC=0.763; p=0.0108) **(Figure 4H)**.

These observations suggest that miR-502-3p and miR-501-3p could be potential serum biomarkers for AD, and ROC curve analysis showed their discriminating power between UC, MCI, and AD.

### 3. Fibroblast study

#### MiR-501-3p and MiR-502-3p Expression Analysis in Fibroblast Samples

In addition to peripheral biomarkers, we were interested in investigating how miR-501-3p and miR-502-3p changed in other cells of the body. As such, we evaluated miR-501-3p and miR-502-3p levels in fibroblasts generated from patients with fAD (n=4), sAD (n=6), and UC (n=8). MiR-501-3p levels were significantly upregulated in fAD relative to UC (p=0.0106); however, they were not significantly increased in sAD relative to UC **(Figure 5A).** MiR-502-3p levels were not significantly different between UC, fAD, and sAD **(Figure 5B).**

**Figure 5.**
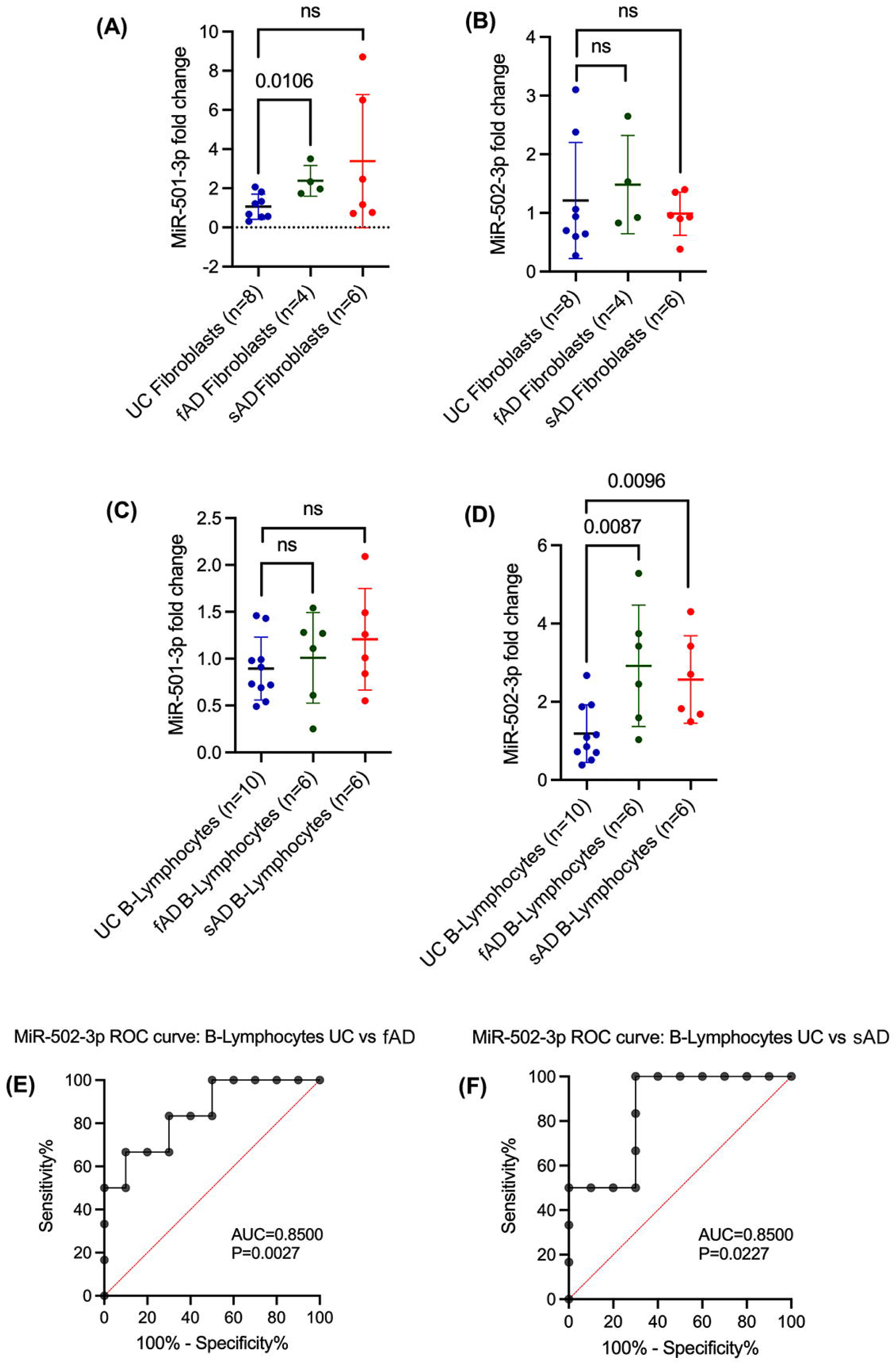
Expression of miR-501-3p and miR-502-3p in UC, fAD, and sAD fibroblasts and B-lymphocytes. **(A)** Expression of miR-501-3p in UC fibroblasts (n=8) compared with fAD fibroblasts (n=4, t-test, p=0.0106) and sAD fibroblasts (n=6, t-test, ns). **(B)** Expression of miR-502-3p in UC fibroblasts (n=8) compared with fAD fibroblasts (n=4, t-test, p=ns) and sAD fibroblasts (n=6, t-test, ns). **(C)** Expression of miR-501-3p in UC B-lymphocytes (n=10) compared with fAD B-lymphocytes (n=6, t-test, ns) and sAD B-lymphocytes (n=6, t-test, ns). **(D)** Expression of miR-502-3p in UC B-lymphocytes (n=10) compared with fAD B-lymphocytes (n=6, t-test, p=0.0087) and sAD B-lymphocytes (n=6, t-test, p=0.0096). **(E)** ROC curve analysis of miR-502-3p expression in UC vs. fAD B-lymphocytes (AUC=0.8500, p=0.0027). **(F)** ROC curve analysis of miR-502-3p expression in UC vs. sAD B-lymphocytes (AUC=0.8500, p=0.0227). ns: not significant.

### 4. B-Lymphocyte study

#### MiR-501-3p and MiR-502-3p Expression in Analysis in B-Lymphocyte Samples

Further, we assessed miR-501-3p and miR-502-3p levels in B-lymphocytes obtained from patients with fAD (n=6), sAD (n=6), and UC (n=10). MiR-501-3p levels were not significantly different between UC, fAD, and sAD **(Figure 5C).** MiR-502-3p levels were significantly upregulated in fAD relative to UC (p=0.0087) and in sAD relative to UC (p=0.0096) **(Figure 5D).** Additionally, we conducted a ROC curve analysis to assess the diagnostic value of miR-502-3p fold change in B-lymphocytes. Our results show a significant AUC for miR-502-3p fold change in fAD vs. UC (AUC=0.8500; p=0.0027) **(Figure 5E)** and sAD vs. UC (AUC=0.8500; p=0.0227) **(Figure 5F)**. The B-lymphocyte data further corroborate the circulatory biomarker nature of miR-502-3p in AD.

### 5. Cells Study

#### APP and Tau Plasmids Expression in Human Neuroblastoma Cells

Finally, we aimed to evaluate how the expression of miR-501-3p and miR-502-3p change intracellularly and if it is secreted extracellularly from the cells in response to AD pathology. First, SH-SY5Y cells were transfected with control, APP, and Tau plasmids. Then, the expressions of APP and Tau proteins were confirmed by immunoblotting and qRT-PCR analysis. Immunoblotting analysis revealed high levels of both APP and Tau proteins relative to control **(Figure 6A)**. Densitometry analysis of APP and Tau protein showed a significant upregulation of APP (p<0.001) and Tau (p=0.0016) in the cells transfected with APP and Tau plasmids, compared to control plasmid-transfected cells **(Figure 6B)**. Subsequently, a significant fold upregulation of APP (∼280-fold; p<0.001) and Tau (∼125-fold; p<0.001) mRNA was observed in the cells transfected with APP and Tau plasmids relative to the control plasmid **(Figure 6C)**.

**Figure 6.**
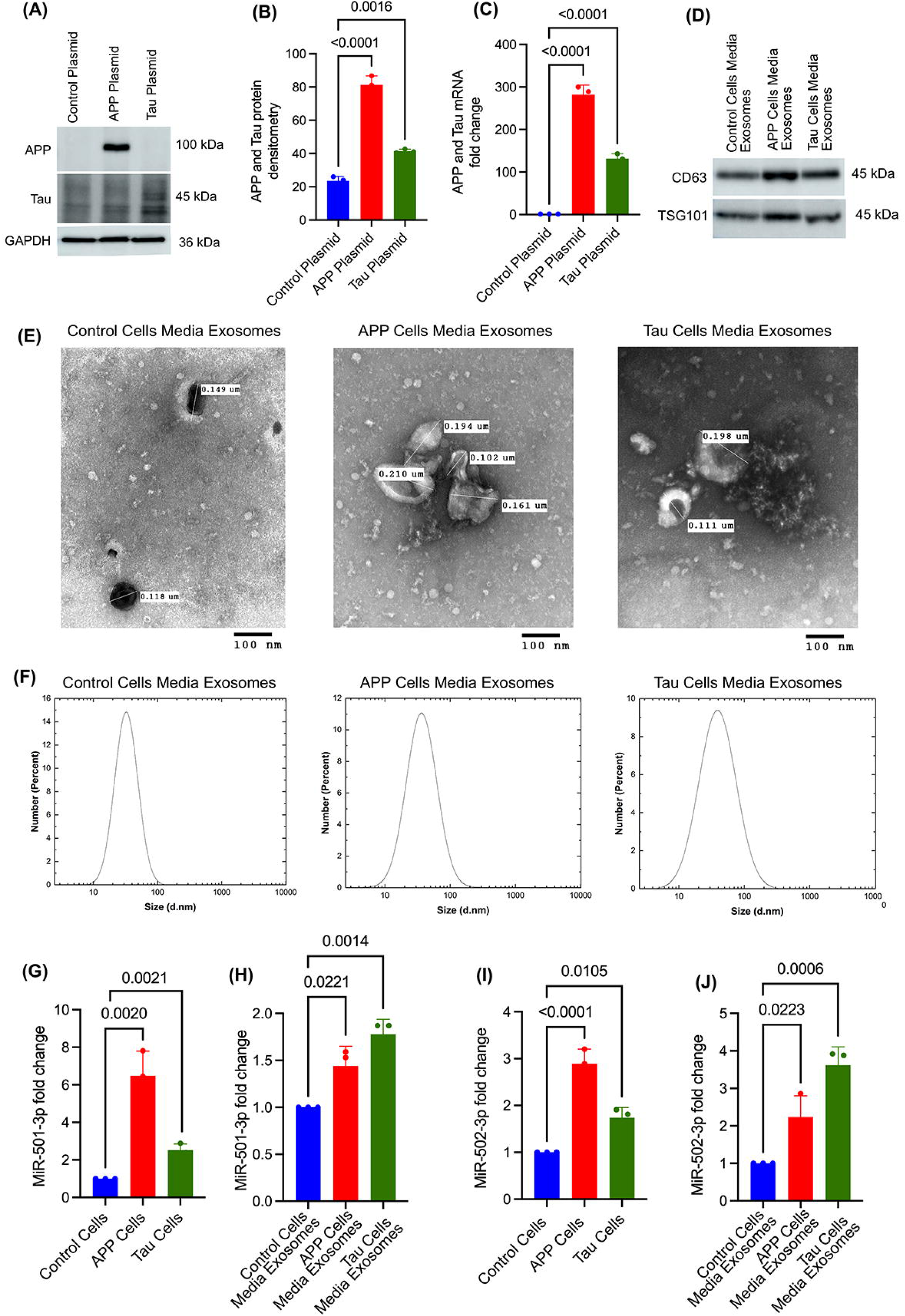
Status of miR-501-3p and miR-502-3p in APP-transfected and Tau-transfected cells and media exosomes. **(A)** Immunoblotting analysis of APP, Tau, and GAPDH proteins. **(B)** APP densitometry analysis in control plasmid-transfected cells vs. APP-transfected cells (t-test, p<0.0001) and Tau densitometry in control plasmid-transfected cells vs. Tau-transfected cells (t-test, p=0.0016). **(C)** APP mRNA fold change in control plasmid-transfected cells vs. APP-transfected cells (t-test, p<0.0001) and Tau mRNA fold change in control plasmid-transfected cells vs. Tau-transfected cells (t-test, p<0.0001). **(D)** Immunoblotting analysis of exosome marker proteins CD63 and TSG101. **(E)** TEM images of exosomes from control plasmid-transfected cell media, APP-transfected cell media, and Tau-transfected cell media at 100 nm magnification. **(F)** Particle size analysis of exosomes in control, APP and Tau-plasmids transfected SH-SY5Y cells media exosomes using DLS. **(G)** Expression of miR-501-3p in control plasmid-transfected cells compared with APP-transfected cells (t-test, p=0.0020) and Tau-transfected cells (t-test, p=0.0021). **(H)** Expression of miR-501-3p in exosomes isolated from control plasmid-transfected cell media compared with APP-transfected cell media (t-test, p=0.0221) and Tau-transfected cell media (t-test, p=0.0014). **(I)** Expression of miR-502-3p in control plasmid-transfected SH-SY5Y cells compared with APP-transfected SH-SY5Y cells (t-test, p<0.0001) and Tau-transfected SH-SY5Y cells (t-test, p=0.0105). **(J)** Expression of miR-502-3p in exosomes isolated from control plasmid-transfected SH-SY5Y cell media compared with APP-transfected SH-SY5Y cell media (t-test, p=0.0223) and Tau-transfected SH-SY5Y cell media (t-test, p=0.0006).

#### Characterization of Cell Media Exosomes

Exosomes were isolated from the media of SH-SY5Y cells expressing the control, APP, and Tau plasmids. The isolated exosomes were characterized by immunoblotting, TEM, and DLS analysis. The immunoblotting analysis showed detectable levels of CD63 and TSG101 proteins in all three groups of media exosomes **(Figure 6D)**. Next, TEM analysis showed the cup-shaped ultrastructure of exosomes in the media from control plasmid, APP-plasmid, and Tau-plasmid transfected cells **(Figure 6E)**. The exosome vesicle size ranged from 0.102 to 0.210 μm in all three groups – control, APP, and Tau cell media exosomes with no significant size differences observed between the groups. Finally, DLS analysis demonstrated exosome size ranging from 10-100 nm, with peak measurements consistently around 40 nm across all three cell media exosome groups **(Figure 6F)**. Further, miR-501-3p and miR-502-3p expressions were analyzed inside the cells and media exosomes.

#### MiR-501-3p Expression Analysis in APP and Tau Transfected Cells and Media Exosomes

To determine the miR-501-3p expression changes intracellularly and extracellularly, qRT-PCR analysis was performed for miR-501-3p on APP– and Tau-expressing cells and their media exosomes. MiR-501-3p level was significantly upregulated in the cells treated with APP (p=0.0020) and Tau (p=0.0021) cells to control cells **(Figure 6G)**. Similarly, elevated expression of miR-501-3p was detected in the media exosomes of APP (p=0.0221) and Tau (p=0.0014) plasmid-treated cells **(Figure 6H)**.

#### MiR-502-3p Expression Analysis in APP and Tau Transfected Cells and Media Exosomes

Likewise, intracellular expression of miR-502-3p was also significantly elevated in APP-(p<0.0001) and Tau-(p=0.0105) expressing cells relative to control cells **(Figure 6I)**. Further, extracellular expression of miR-502-3p was also elevated in the media exosomes of APP-(p=0.023) and Tau-(p=0.0006) expressing cells relative to control media exosomes **(Figure 6J)**. These observations confirm that miR-501-3p and miR-502-3p expressions change inside the cells, and they are secreted out from the cells in response to both APP and Tau pathology.

## DISCUSSION

Diagnostic methods to accurately detect AD remain limited, and similarly, not many treatment modalities currently exist to address AD. With the recent advent of aducanumab, a drug that slows disease progression, there is an increasing need to discover more effective ways to detect AD. In 2011, the National Institute of Neurological and Communicative Disorders and Stroke (NINCDS) and the Alzheimer’s Disease and Related Disorders Association (ADRDA) revised the diagnostic criteria for AD to include Aβ40, Aβ42, Aβ42/Aβ40, t-tau, and p-tau as CSF biomarkers.^38,39^ However, these biomarkers still struggle to detect the early stages of AD, highlighting the need for continued research to identify new reliable biomarkers. The first established miRNA biomarker was used for diffuse large B-cell lymphoma in 2008.^40^ Since then, the biomarker potential of miRNAs has been heavily studied in a multitude of diseases, including AD.^41–44^ While several potential miRNA biomarkers have been identified for AD, none have yet been established for clinical use. With this study, we continue to expand the list of possible miRNA biomarkers and characterize their clinical potential. We have identified increased expressions of miR-501-3p and miR-502-3p in the AD synapses and have correlated their expression with disease progression based on Braak staging.^32,33^ Moreover, our lab is investigating the role of miR-502-3p in the modulation of GABAergic synapse function in AD.^34,45^ Therefore, we are interested in evaluating their diagnostic biomarker potential. Many existing studies explored miRNAs in only one sample type (CSF, serum, etc.), which limits our understanding of how miRNAs change in the body during AD pathology. In this study, we analyzed miR-501-3p and miR-502-3p status in AD CSF exosomes, AD serum exosomes, AD fibroblasts, B-lymphocytes, AD cells and media exosomes.

First, we studied the status of miR-501-3p and miR-502-3p in the AD CSF exosomes. Since CSF is in direct contact with the brain, it is possible that brain cells secrete exosomes enriched with various biomolecules (proteins, miRNAs, metabolites, etc.) in response to AD or neurodegeneration.^46^ Moreover, CSF contains a wide range of biomarkers that can indicate the presence of various neurological diseases.^47^ In AD, for example, CSF levels of amyloid-beta (Aβ), tau proteins, and phosphorylated tau have been shown to correlate with disease stages.^48^ Exosomes contain miRNAs, proteins, and other nucleic acids, that are packaged within exosomes, which are secreted by various cell types and commonly found in both CSF and serum.^49^ To identify the sources of these exosomes, we performed immunoblotting for cell type-specific markers. Immunoblotting analysis revealed that most exosomes secreted in the CSF originate from neurons and astrocytes. Our qRT-PCR results showed a significant increase in the levels of both miR-501-3p and miR-502-3p in the AD CSF samples compared to UC CSF samples. These findings are consistent with the overexpression of these miRNAs observed in AD synaptosomes.^32^ Since these miRNAs are elevated at the synapse,^32^ and neurons and astrocytes are major players in synapse formation,^50^ it is possible that defective synapses in AD may cause the exosome-mediated heavy secretion of these miRNAs into the CSF. Moreover, significant AUC values from ROC curve analyses further support their biomarker capabilities. Given the clear differential expression of miR-501-3p and miR-502-3p, a potential threshold could be established to support AD diagnosis. Next, we compared miRNA fold change with the neuropathologic reports of the CSF samples to evaluate whether miRNA levels in the CSF are associated with the severity of amyloid plaques and NFT found in affected brain regions in AD. Upon reviewing the neuropathology reports, we observed that the entorhinal cortex exhibited the highest number of samples with severe amyloid plaques and NFT, followed by the amygdala and hippocampus. Interestingly, most hippocampus or amygdala samples did not show greater severity than the entorhinal cortex areas. This observation suggests that AD pathology progresses in a predictable pattern, with the entorhinal cortex being the first affected brain area in AD.^51^ As mentioned previously, a quantitative analysis of our data is challenging due to the uneven distribution of samples across different severity categories. However, preliminary observations suggest that differential miR-501-3p and miR-502-3p fold changes could reflect the severity of AD pathology in specific brain regions. For example, we found a potential correlation between miR-501-3p fold changes and amyloid plaque severity in the entorhinal cortex and between miR-502-3p fold changes and NFT severity in the amygdala. To our knowledge, no studies explored the relationship between miRNA fold changes and the neuropathological severity of AD. This is an area we are particularly interested in investigating further, as such a study could contribute to earlier diagnosis of AD based on miRNA levels.

To further assess the biomarker potential of miR-501-3p and miR-502-3p, we next evaluated their status in AD, MCI, and control serum samples. In the serum, we found that the exosomes could potentially be derived from neurons, astrocytes, and microglia. Since serum is more heterogenous than CSF, it could be possible that exosomes may be secreted from a wide range of cells and tissues affected in AD. Our qRT-PCR results reveal upregulation of both miR-501-3p and miR-502-3p in AD and MCI serum exosomes. The gradual upregulation of these miRNAs in control vs. MCI vs. AD further supports the early AD detection capabilities of these miRNAs. Our findings are well aligned with previous studies of miR-501-3p.^52,53^ Toyama et al., unveiled the elevated levels of miR-501-3p on blood exosomes in MCI patients relative to controls.^53^ Hara et al. reported the upregulation of miR-501-3p in AD brain. However, they found reduced levels of miR-501-3p in the serum.^52^ One reason for this discrepancy could be that they studied miR-501-3p levels directly from the serum and not the exosomes. Another possibility could be that they used serum samples collected within two weeks of the patient’s deaths, while our samples were collected much earlier before their deaths. Exosome biogenesis and subsequent release require energy,^54,55^ and advanced age or severe AD may lead to a decline in ATP levels, potentially affecting exosome production and miRNA levels in the serum.^56,57^ Further studies must be done to explore how serum miRNA levels change with age or with the severity of diseases. Altogether, our findings support the potential of miR-501-3p and miR-502-3p as serum biomarkers for AD.

Additionally, we investigated the status of miR-501-3p and miR-502-3p in other peripheral cells, such as fibroblasts and B-lymphocytes. The cellular and molecular functions of fibroblasts and B-lymphocytes are dysregulated during AD pathogenesis.^58,59^ Our results show that miR-501-3p is upregulated in fAD fibroblasts; however, there was no significant difference in miR-501-3p in sAD fibroblasts. Additionally, there was no significant difference in miR-501-3p levels in fAD or sAD B-lymphocytes. Interestingly, miR-502-3p showed an inverse pattern: it was upregulated in both fAD and sAD B-lymphocytes, but no significant difference in miR-502-3p expression was observed in fAD or sAD fibroblasts. While further studies are needed to determine whether miRNA levels in fibroblasts or B-lymphocytes could serve as reliable biomarkers, our results suggest that miR-501-3p may be associated with fibroblasts and miR-502-3p could be associated with B-lymphocytes dysfunction during AD pathology.

Lastly, we investigated how AD pathology might directly influence miR-501-3p and miR-502-3p levels in intracellular and extracellular secretions. We found that both miRNAs were upregulated intracellularly and extracellularly in the cells overexpressing APP and tau proteins. In APP overexpressed cells, miR-501-3p and miR-502-3p fold expressions were higher intracellularly than extracellularly. In contrast, tau overexpressed cells exhibited a higher fold change of miR-502-3p in the extracellular media exosomes than intracellularly. This suggests that tau-related AD pathology may be directly associated with the peripheral secretion of miR-502-3p.

Further studies with larger sample sizes, including patients with different ethnic groups, are warranted before the preclinical testing of these miRNAs.

In conclusion, our study thoroughly investigated the biomarker potential of miR-501-3p and miR-502-3p across different types of AD samples. Altogether, our current findings help us to conclude that miR-501-3p and miR-502-3p could be a potential biomarker panel and a representation of AD pathology. While we could not confidently define their clinical diagnostic value, the current study has laid a foundation for future research on miR-501-3p and miR-502-3p and their potential roles in predicting early AD.

## ACKNOWLEDGMENTS

We are thankful to the NIH NeuroBioBanks for providing the AD and control CSF specimens. We are thankful to the Coriell Institute of Medical Research, for providing the AD and control subjects’ Fibroblasts and B-Lymphocytes for our study. We are thankful to the Texas Alzheimer’s Research and Care Consortium for proving the AD, MCI and control serum samples. We are also thankful to the Dr. Hamidreza Sharifan, Dr. Md. Nurunnabi, Ms. Kavitha Beluri and Ms. Anusha Reddy, University of Texas El Paso for helping the particle size analysis. We are exceedingly grateful to Prof. Rajkumar Lakshmanaswamy, Chair of the Department of Molecular and Translational Medicine, TTUHSC El Paso for the immense research support. We would also like to thank our lab members Ms. Melissa Torres, Mrs. Sheryl Rodriguez, Ms. Yogyana Santana-Salais and Ms. Angelica Amaya.

## CONFLICT OF INTEREST

The author would like to inform that he filed a patent on “Synaptosomal miRNAs and Synapse Functions in Alzheimer’s Disease” TTU Ref. No. 2022-016, U.S. Patent App. No. PCT/US2023/019298 on Oct 26, 2023 related to the contents of this manuscript. The other authors declare that they have no conflict of interest.

## FUNDING SOURCES

This research was funded by the National Institute on Aging (NIA), National Institutes of Health (NIH), grant number K99AG065645, R00AG065645, R00AG065645-04S1, SARP mini grants TTUHSC EP, Edward N. & Margaret G. Marsh Foundation and TTUHSC EP MTM Startup Funds to S.K.

## CONSENT STATEMENT

Consent was not necessary. The samples obtained from the bio banks are operated under their institution’s IRB approval, and they obtained written informed consent from the donors.

## AUTHORS CONTRIBUTION

Conceptualization and supervision: SK; experimental performance: DD, BS, GG, DR, AK, NT and AP; analysis, interpretation, and validation of data: SK, DD, BS and AK; writing and original draft preparation: DD, BS and SK; review, editing, and finalization of manuscript: DD, BS, DR, GG and SK. All authors have read and agreed to the published version of the manuscript.

## DATA AVAILABILITY

Data will be made available on request.

## Notes

### Competing Interest Statement

The author would like to inform that he filed a patent on Synaptosomal miRNAs and Synapse Functions in Alzheimers Disease TTU Ref. No. 2022-016, U.S. Patent App. No. PCT/US2023/019298 on Oct 26, 2023 related to the contents of this manuscript. The other authors declare that they have no conflict of interest.

