## Supplementary material for "MiRNA-501-3p and MiRNA-502-3p: A Promising Biomarker Panel for Alzheimer’s Disease": SI Table 1

**Supplementary Table 1- Demographic and clinical details of CSF samples.**

| **Sample** | **Age** | **Manner of Death** | **Sex** | **Race** | **PMI (hrs)** | **Braak Stage** | **Clinical Brain Diagnosis** |
| --- | --- | --- | --- | --- | --- | --- | --- |
| UC1 | 61 | Natural | F | White | 20 | 0 | No clinical brain diagnosis found |
| UC2 | 60 | Natural | M | White | 20.42 | 1 | No clinical brain diagnosis found |
| UC3 | 55 | Natural | M | Not reported | 16.55 | 0 | No clinical brain diagnosis found |
| UC4 | 62 | Natural | F | Black or African-American | 7 | 0 | No clinical brain diagnosis found |
| UC5 | 56 | Natural | M | Black or African-American | 15.43 | 1 | No clinical brain diagnosis found |
| UC6 | 62 | Natural | F | Not reported | 6.7 | 1 | No clinical brain diagnosis found |
| UC7 | 61 | Natural | M | White | 14.5 | 0 | No clinical brain diagnosis found |
| UC8 | 61 | Natural | M | White | 14.5 | 0 | No clinical brain diagnosis found |
| UC9 | 58 | Natural | F | Black or African-American | 22.58 | 0 | No clinical brain diagnosis found |
| UC10 | 63 | Natural | F | White | 19.52 | 1 | No clinical brain diagnosis found |
| AD1 | 64 | Natural | F | White | 7.5 | 6 | Alzheimer's disease with late onset |
| AD2 | 65 | Natural | F | White | 5.25 | 6 | Alzheimer's disease with late onset |
| AD3 | 61 | Natural | F | White | 6 | 6 | Alzheimer's disease with late onset |
| AD4 | 68 | Natural | M | White | 24.7 | 0 | Alzheimer's disease with early onset |
| AD5 | 67 | Natural | M | White | 6.25 | 6 | Alzheimer's disease with late onset |
| AD6 | 66 | Natural | F | White | 26.25 | 0 | Alzheimer's disease with late onset |
| AD7 | 67 | Natural | M | White | 8.25 | 6 | Alzheimer's disease with late onset |
| AD8 | 69 | Natural | F | Not reported | 8.33 | 6 | Alzheimer's disease with late onset |
| AD9 | 65 | Natural | M | White | 6.22 | 6 | Alzheimer's disease with early onset |
| AD10 | 68 | Natural | M | White | 17.92 | 6 | Alzheimer's disease with late onset |
| AD11 | 62 | Natural | M | White | 7.33 | 6 | Alzheimer's disease with late onset |
| AD12 | 62 | Natural | M | White | 16.48 | 6 | Alzheimer's disease with early onset |
| AD13 | 67 | Natural | M | White | 5.67 | 6 | Alzheimer's disease with late onset |
| AD14 | 57 | Natural | M | White | 9.83 | 0 | Alzheimer's disease with late onset |
| AD15 | 69 | Natural | F | White | 6.5 | 6 | Alzheimer's disease with late onset |
| AD16 | 62 | Natural | M | White | 5.92 | 3 | Alzheimer's disease with late onset |
| AD17 | 62 | Natural | F | White | 3.67 | 6 | Alzheimer's disease with early onset |
| AD18 | 58 | Natural | M | White | 5.83 | 6 | Alzheimer's disease with late onset |
| AD19 | 66 | Natural | F | White | 5.33 | 6 | Alzheimer's disease with late onset |
| AD20 | 65 | Natural | F | White | 4.08 | 6 | Alzheimer's disease with late onset |
| AD21 | 63 | Natural | F | Not reported | 9.42 | 6 | Alzheimer's disease with early onset |
| AD22 | 52 | Natural | F | White | 63 | 6 | Alzheimer's disease with late onset |
| AD23 | 63 | Natural | M | White | 5.83 | 6 | Alzheimer's disease with late onset |
| AD24 | 65 | Natural | F | White | 5.83 | 6 | Alzheimer's disease with late onset |
| AD25 | 68 | Natural | F | White | 6 | 2 | Alzheimer's disease with late onset |
| AD26 | 68 | Natural | M | Black or African-American | 16.47 | 6 | Alzheimer's disease with early onset |
