## Supplementary material for "MiRNA-501-3p and MiRNA-502-3p: A Promising Biomarker Panel for Alzheimer’s Disease": SI Table 2

**Supplementary Table 2- Amyloid plaques and NFT density in the different brain regions of CSF samples.**

| **Sample** | **Clinical Brain Diagnosis** | **Braak Stage** | **Amyloid Plaques - Hippocampus** | **Amyloid Plaques - Amygdala** | **Amyloid Plaques - Entorhinal Cortex** | **NFT - Hippocampus** | **NFT - Amygdala** | **NFT - Entorhinal Cortex** | **Amyloid Plaques - Middle Frontal Gyrus** | **Amyloid Plaques – Inferior Parietal Lobule** | **NFT- Middle Frontal Gyrus** | **NFT- Superior Temporal Gyrus** |
| --- | --- | --- | --- | --- | --- | --- | --- | --- | --- | --- | --- | --- |
| HC1 | No clinical brain diagnosis found | 0 | 0- None | 0- None | 0- None | 0- None | 0- None | 0- None | 0- None | 0- None | 0- None | 0- None |
| HC2 | No clinical brain diagnosis found | 1 | 0- None | 0- None | 0- None | 0- None | 0- None | 0- None | 0- None | 0- None | 0- None | 0- None |
| HC3 | No clinical brain diagnosis found | 0 | 0- None | 0- None | 0- None | 0- None | 0- None | 1- Sparse | 0- None | 0- None | 0- None | 0- None |
| HC4 | No clinical brain diagnosis found | 0 | 0- None | 0- None | 0- None | 0- None | 0- None | 0- None | 0- None | 0- None | 0- None | 0- None |
| HC5 | No clinical brain diagnosis found | 1 | 0- None | 0- None | 0- None | 1- Sparse | 0- None | 1- Sparse | 0- None | 0- None | 0- None | 0- None |
| HC6 | No clinical brain diagnosis found | 1 | 0- None | 0- None | 0- None | 1- Sparse | 0- None | 1- Sparse | 0- None | 0- None | 0- None | 0- None |
| HC7 | No clinical brain diagnosis found | 0 | 0- None | 0- None | 0- None | 0- None | 0- None | 0- None | 0- None | 0- None | 0- None | 0- None |
| HC8 | No clinical brain diagnosis found | 0 | 0- None | 0- None | 0- None | 0- None | 0- None | 0- None | 0- None | 0- None | 0- None | 0- None |
| HC9 | No clinical brain diagnosis found | 0 | 0- None | 0- None | 0- None | 0- None | 0- None | 1- Sparse | 0- None | 0- None | 0- None | 0- None |
| HC10 | No clinical brain diagnosis found | 1 | 0- None | 0- None | 0- None | 1- Sparse | 0- None | 1- Sparse | 0- None | 1- Sparse | 0- None | 0- None |
| AD1 | Alzheimer's disease with late onset | 6 | 1- Sparse | 5- Frequent/Severe | 5- Frequent/Severe | 5- Frequent/Severe | 5- Frequent/Severe | 5- Frequent/Severe | 5- Frequent/Severe | 5- Frequent/Severe | 5- Frequent/Severe | 5- Frequent/Severe |
| AD2 | Alzheimer's disease with late onset | 6 | 0- None | 5- Frequent/Severe | 5- Frequent/Severe | 3- Moderate | 5- Frequent/Severe | 5- Frequent/Severe | 5- Frequent/Severe | 5- Frequent/Severe | 3- Moderate | 3- Moderate |
| AD3 | Alzheimer's disease with late onset | 6 | 3- Moderate | 9- NA | 3- Moderate | 3- Moderate | 9- NA | 5- Frequent/Severe | 5- Frequent/Severe | 5- Frequent/Severe | 5- Frequent/Severe | 3- Moderate |
| AD4 | Alzheimer's disease with early onset | 0 | 0- None | 1- Sparse | 0- None | 0- None | 0- None | 0- None | 1- Sparse | 0- None | 0- None | 0- None |
| AD5 | Alzheimer's disease with late onset | 6 | 5- Frequent/Severe | 3- Moderate | 5- Frequent/Severe | 5- Frequent/Severe | 5- Frequent/Severe | 5- Frequent/Severe | 5- Frequent/Severe | 5- Frequent/Severe | 5- Frequent/Severe | 5- Frequent/Severe |
| AD6 | Alzheimer's disease with late onset | 0 | 0- None | 0- None | 0- None | 0- None | 0- None | 1- Sparse | 0- None | 0- None | 0- None | 0- None |
| AD7 | Alzheimer's disease with late onset | 6 | 3- Moderate | 5- Frequent/Severe | 3- Moderate | 5- Frequent/Severe | 5- Frequent/Severe | 5- Frequent/Severe | 5- Frequent/Severe | 3- Moderate | 5- Frequent/Severe | 5- Frequent/Severe |
| AD8 | Alzheimer's disease with late onset | 6 | 1- Sparse | 5- Frequent/Severe | 1- Sparse | 5- Frequent/Severe | 3- Moderate | 5- Frequent/Severe | 5- Frequent/Severe | 3- Moderate | 5- Frequent/Severe | 5- Frequent/Severe |
| AD9 | Alzheimer's disease with early onset | 6 | 1- Sparse | 1- Sparse | 1- Sparse | 5- Frequent/Severe | 5- Frequent/Severe | 5- Frequent/Severe | 5- Frequent/Severe | 3- Moderate | 5- Frequent/Severe | 5- Frequent/Severe |
| AD10 | Alzheimer's disease with late onset | 6 | 3- Moderate | 5- Frequent/Severe | 5- Frequent/Severe | 5- Frequent/Severe | 3- Moderate | 5- Frequent/Severe | 5- Frequent/Severe | 5- Frequent/Severe | 5- Frequent/Severe | 5- Frequent/Severe |
| AD11 | Alzheimer's disease with late onset | 6 | 3- Moderate | 5- Frequent/Severe | 5- Frequent/Severe | 3- Moderate | 5- Frequent/Severe | 5- Frequent/Severe | 5- Frequent/Severe | 5- Frequent/Severe | 5- Frequent/Severe | 5- Frequent/Severe |
| AD12 | Alzheimer's disease with early onset | 6 | 1- Sparse | 3- Moderate | 5- Frequent/Severe | 5- Frequent/Severe | 1- Sparse | 3- Moderate | 5- Frequent/Severe | 5- Frequent/Severe | 5- Frequent/Severe | 5- Frequent/Severe |
| AD13 | Alzheimer's disease with late onset | 6 | 3- Moderate | 3- Moderate | 5- Frequent/Severe | 5- Frequent/Severe | 3- Moderate | 5- Frequent/Severe | 5- Frequent/Severe | 5- Frequent/Severe | 5- Frequent/Severe | 5- Frequent/Severe |
| AD14 | Alzheimer's disease with late onset | 0 | 0- None | 0- None | 0- None | 0- None | 0- None | 1- Sparse | 0- None | 0- None | 0- None | 0- None |
| AD15 | Alzheimer's disease with late onset | 6 | 1- Sparse | 3- Moderate | 5- Frequent/Severe | 5- Frequent/Severe | 3- Moderate | 5- Frequent/Severe | 5- Frequent/Severe | 5- Frequent/Severe | 5- Frequent/Severe | 5- Frequent/Severe |
| AD16 | Alzheimer's disease with late onset | 3 | 0- None | 5- Frequent/Severe | 0- None | 3- Moderate | 1- Sparse | 3- Moderate | 5- Frequent/Severe | 5- Frequent/Severe | 0- None | 0- None |
| AD17 | Alzheimer's disease with early onset | 6 | 1- Sparse | 3- Moderate | 1- Sparse | 1- Sparse | 0- None | 5- Frequent/Severe | 5- Frequent/Severe | 5- Frequent/Severe | 3- Moderate | 3- Moderate |
| AD18 | Alzheimer's disease with late onset | 6 | 3- Moderate | 3- Moderate | 3- Moderate | 5- Frequent/Severe | 3- Moderate | 5- Frequent/Severe | 3- Moderate | 3- Moderate | 3- Moderate | 3- Moderate |
| AD19 | Alzheimer's disease with late onset | 6 | 3- Moderate | 3- Moderate | 3- Moderate | 5- Frequent/Severe | 3- Moderate | 5- Frequent/Severe | 5- Frequent/Severe | 3- Moderate | 5- Frequent/Severe | 5- Frequent/Severe |
| AD20 | Alzheimer's disease with late onset | 6 | 3- Moderate | 5- Frequent/Severe | 3- Moderate | 5- Frequent/Severe | 5- Frequent/Severe | 5- Frequent/Severe | 5- Frequent/Severe | 5- Frequent/Severe | 5- Frequent/Severe | 5- Frequent/Severe |
| AD21 | Alzheimer's disease with early onset | 6 | 5- Frequent/Severe | 5- Frequent/Severe | 3- Moderate | 5- Frequent/Severe | 5- Frequent/Severe | 5- Frequent/Severe | 3- Moderate | 5- Frequent/Severe | 5- Frequent/Severe | 5- Frequent/Severe |
| AD22 | Alzheimer's disease with late onset | 6 | 5- Frequent/Severe | 5- Frequent/Severe | 5- Frequent/Severe | 5- Frequent/Severe | 3- Moderate | 5- Frequent/Severe | 5- Frequent/Severe | 5- Frequent/Severe | 5- Frequent/Severe | 3- Moderate |
| AD23 | Alzheimer's disease with late onset | 6 | 3- Moderate | 3- Moderate | 5- Frequent/Severe | 5- Frequent/Severe | 1- Sparse | 5- Frequent/Severe | 5- Frequent/Severe | 5- Frequent/Severe | 5- Frequent/Severe | 5- Frequent/Severe |
| AD24 | Alzheimer's disease with late onset | 6 | 1- Sparse | 3- Moderate | 5- Frequent/Severe | 5- Frequent/Severe | 3- Moderate | 5- Frequent/Severe | 5- Frequent/Severe | 5- Frequent/Severe | 5- Frequent/Severe | 5- Frequent/Severe |
| AD25 | Alzheimer's disease with late onset | 2 | 1- Sparse | 3- Moderate | 1- Sparse | 1- Sparse | 0- None | 3- Moderate | 1- Sparse | 3- Moderate | 0- None | 0- None |
| AD26 | Alzheimer's disease with early onset | 6 | 1- Sparse | 3- Moderate | 3- Moderate | 5- Frequent/Severe | 3- Moderate | 3- Moderate | 3- Moderate | 3- Moderate | 5- Frequent/Severe | 5- Frequent/Severe |
