## Supplementary material for "MiRNA-501-3p and MiRNA-502-3p: A Promising Biomarker Panel for Alzheimer’s Disease": SI Table 3

**Supplementary Table 3- Demographic and clinical details of serum samples.**

| **Sample** | **Age** | **Sex** | **Race** | **MMSE** | **APOE Genotype** | **Total Tau (pg/ml)** |
| --- | --- | --- | --- | --- | --- | --- |
| AD1 | 82 | M | White | 22 | e3/e4 | 0.5724222 |
| AD2 | 85 | F | Black or African-American | 20 | e3/e4 | 0.1593416 |
| AD3 | 71 | M | White | 29 | e3/e4 | NA |
| AD4 | 79 | F | White | 11 | e3/e4 | 0.1495421 |
| AD5 | 81 | M | White | 16 | e3/e3 | 0.1074443 |
| AD6 | 75 | M | White | 24 | e3/e4 | 0.221794 |
| AD7 | 81 | F | White | 27 | e3/e3 | NA |
| AD8 | 79 | F | White | 19 | e3/e3 | 0.3239219 |
| AD9 | 84 | M | N/A | 24 | NA | NA |
| AD10 | 82 | M | N/A | 22 | NA | NA |
| AD11 | 88 | M | N/A | 19 | NA | NA |
| AD12 | 76 | F | N/A | 24 | NA | NA |
| AD13 | 75 | F | N/A | 28 | NA | NA |
| AD14 | 71 | M | N/A | 23 | NA | NA |
| AD15 | 76 | F | N/A | 23 | NA | NA |
| AD16 | 67 | F | White | 25 | NA | NA |
| AD17 | 68 | M | N/A | 18 | NA | NA |
| AD18 | 72 | M | N/A | 24 | NA | NA |
| AD19 | 75 | M | N/A | 25 | NA | NA |
| AD20 | 81 | F | N/A | 11 | NA | NA |
| AD21 | 58 | F | N/A | 22 | NA | NA |
| AD22 | 66 | M | N/A | 28 | NA | NA |
| AD23 | 63 | F | N/A | 17 | NA | NA |
| AD24 | 81 | F | White | 10 | NA | NA |
| AD25 | 70 | F | Black or African-American | 10 | NA | NA |
| MCI1 | 86 | F | White | 16 | e3/e3 | 0.6021046 |
| MCI2 | 68 | F | White | 16 | e3/e3 | 0.4391071 |
| MCI3 | 70 | M | Asian | 18 | e3/e3 | NA |
| MCI4 | 77 | F | White | 13 | e3/e3 | 0.2722061 |
| MCI5 | 75 | M | White | 17 | e3/e4 | 0.3603593 |
| MCI6 | 71 | F | N/A | 17 | NA | NA |
| MCI7 | 64 | F | N/A | 19 | NA | NA |
| MCI8 | 59 | F | N/A | 17 | NA | NA |
| MCI9 | 71 | F | N/A | 17 | NA | NA |
| MCI10 | 66 | M | N/A | 19 | NA | NA |
| MCI11 | 67 | M | N/A | 19 | NA | NA |
| MCI12 | 53 | F | N/A | 17 | NA | NA |
| MCI13 | 72 | F | N/A | 15 | NA | NA |
| MCI14 | 69 | M | N/A | 16 | NA | NA |
| MCI15 | 77 | F | N/A | 18 | NA | NA |
| UC1 | 62 | F | White | 29 | e3/e3 | NA |
| UC2 | 52 | M | White | 28 | e3/e3 | 0.2426996 |
| UC3 | 87 | F | N/A | 28 | NA | NA |
| UC4 | 81 | F | White | 28 | NA | NA |
| UC5 | 76 | F | N/A | 28 | NA | NA |
| UC6 | 73 | F | N/A | 30 | NA | NA |
| UC7 | 70 | F | N/A | 28 | NA | NA |
| UC8 | 61 | F | N/A | 20 | NA | NA |
| UC9 | 62 | F | N/A | 30 | NA | NA |
| UC10 | 62 | M | N/A | 29 | NA | NA |
| UC11 | 72 | M | N/A | 21 | NA | NA |
| UC12 | 65 | M | N/A | 30 | NA | NA |
| UC13 | 65 | M | N/A | 30 | NA | NA |
| UC14 | 53 | F | N/A | 30 | NA | NA |
| UC15 | 72 | M | N/A | 29 | NA | NA |
| UC16 | 83 | F | N/A | 29 | NA | NA |
| UC17 | 75 | F | N/A | 30 | NA | NA |
| UC18 | 62 | F | N/A | 28 | NA | NA |
| UC19 | 66 | F | N/A | 20 | NA | NA |

NA* Not available
