## Supplementary material for "MiRNA-501-3p and MiRNA-502-3p: A Promising Biomarker Panel for Alzheimer’s Disease": SI Table 4

**Supplementary Table 4- Demographic detail of fibroblasts samples.**

| **Serial Number** | **Catalog Number** | **Passage Number** | **Sex** | **Age** | **Biopsy Sources** | **Tissue Type** | **Race** | **Disease Status** |
| --- | --- | --- | --- | --- | --- | --- | --- | --- |
| 1 | AG02261 | 11 | M | 61 | Abdomen | Skin | Caucasian | Unaffected control |
| 2 | AG16104 | 6 | F | 55 | Arm | Skin | Black | Unaffected control |
| 3 | AG16086 | 6 | F | 67 | Arm | Skin | Other | Unaffected control |
| 4 | AG12207 | 13 | M | 68 | Arm | Skin | N/A | Unaffected control |
| 5 | AG02258 | 6 | F | 46 | Lung | Lung | Caucasian | Unaffected control |
| 6 | AG02262 | 4 | M | 61 | Lung | Lung | Caucasian | Unaffected control |
| 7 | AG06561 | 5 | F | 16FW (Fetal week) | Sacrum | Skin | Caucasian | Unaffected control |
| 8 | AG12211 | 11 | M | 54 | Lung | Lung | Caucasian | Unaffected control |
| 9 | AG05810 | 11 | F | 79 | Arm | Skin | Caucasian | Familial AD |
| 10 | AG06844 | 12 | M | 59 | Arm | Skin | Caucasian | Familial AD |
| 11 | AG07613 | 16 | M | 66 | Arm | Skin | Caucasian | Familial AD |
| 12 | AG09908 | 14 | F | 81 | Arm | Skin | Caucasian | Familial AD |
| 13 | AG04400 | 19 | F | 61 | Skin | Skin | Caucasian | Sporadic AD |
| 14 | AG06263 | 11 | F | 67 | Arm | Skin | Caucasian | Sporadic AD |
| 15 | AG06264 | 7 | F | 62 | Arm | Skin | N/A | Sporadic AD |
| 16 | AG07375 | 6 | M | 71 | Arm | Skin | Caucasian | Sporadic AD |
| 17 | AG08243 | 7 | M | 72 | Arm | Skin | Caucasian | Sporadic AD |
| 18 | AG11368 | 15 | M | 77 | Skin | Skin | Caucasian | Sporadic AD |
