## Supplementary material for "MiRNA-501-3p and MiRNA-502-3p: A Promising Biomarker Panel for Alzheimer’s Disease": SI Table 5

**Supplementary Table 5- Demographic details of B-lymphocytes samples.**

| **Serial Number** | **Catalog Number** | **Sex** | **Age** | **Biopsy Sources** | **Tissue Type** | **Race** | **Disease Status** |
| --- | --- | --- | --- | --- | --- | --- | --- |
| 1 | AG16639 | M | 77 | Peripheral vein | Blood | Caucasian | Unaffected control |
| 2 | AG11684 | M | 82 | Peripheral vein | Blood | Caucasian | Unaffected control |
| 3 | AG12034 | F | 80 | Peripheral vein | Blood | Caucasian | Unaffected control |
| 4 | AG11716 | M | 98 | Peripheral vein | Blood | Caucasian | Unaffected control |
| 5 | AG12032 | M | 84 | Peripheral vein | Blood | Caucasian | Unaffected control |
| 6 | AG16804 | F | 90 | Peripheral vein | Blood | Caucasian | Unaffected control |
| 7 | AG16927 | M | 85 | Peripheral vein | Blood | Caucasian | Unaffected control |
| 8 | AG16973 | F | 80 | Peripheral vein | Blood | Caucasian | Unaffected control |
| 9 | AG10673 | F | 85 | Peripheral vein | Blood | Black | Unaffected control |
| 10 | AG16907 | F | 88 | Peripheral vein | Blood | Caucasian | Unaffected control |
| 11 | AG08242 | M | 72 | Peripheral vein | Blood | Caucasian | Familial AD |
| 12 | AG09905 | M | 72 | Peripheral vein | Blood | Caucasian | Familial AD |
| 13 | AG09907 | F | 71 | Peripheral vein | Blood | Caucasian | Familial AD |
| 14 | AG11755 | F | 85 | Peripheral vein | Blood | Caucasian | Familial AD |
| 15 | AG11757 | F | 81 | Peripheral vein | Blood | Caucasian | Familial AD |
| 16 | AG11758 | M | 83 | Peripheral vein | Blood | Caucasian | Familial AD |
| 17 | AG06204 | M | 67 | Peripheral vein | Blood | Caucasian | Sporadic AD |
| 18 | AG06868 | F | 60 | Peripheral vein | Blood | Caucasian | Sporadic AD |
| 19 | AG11366 | M | 52 | Peripheral vein | Blood | Caucasian | Sporadic AD |
| 20 | AG17512 | M | 70 | Peripheral vein | Blood | African-American | Sporadic AD |
| 21 | AG17529 | F | 86 | Peripheral vein | Blood | African-American | Sporadic AD |
| 22 | AG17574 | F | 83 | Peripheral vein | Blood | African-American | Sporadic AD |
