## Supplementary material for "MiRNA-501-3p and MiRNA-502-3p: A Promising Biomarker Panel for Alzheimer’s Disease": SI Table 6

**Supplementary Table 6- Summary of antibody dilutions and conditions used in the immunoblotting analysis**

| **Marker(s)** | **Primary Antibody and Dilution(s)**  **(4^°^C, overnight)** | **Purchased from Company, City & State** | **Secondary Antibody, Dilution(s)**  **(Room temperature, 2 h)** | **Purchased from Company, City & State** |
| --- | --- | --- | --- | --- |
| CD9  (60232-1-Ig) | Mouse monoclonal  1:500 | Proteintech, Rosemont, IL | Rabbit anti-mouse IgG HRP 1:10,000  (A9044-2 mL) | Millipore Sigma  Burlington, MA |
| CD63  (67605-1-Ig) | Mouse monoclonal  1:500 | Proteintech, Rosemont, IL | Rabbit anti-mouse IgG HRP 1:10,000  (A9044-2 mL) | Millipore Sigma  Burlington, MA |
| TSG101  (NB200-112) | Mouse monoclonal  1:500 | Novus Biologicals, Minneapolis, MN | Rabbit anti-mouse IgG HRP 1:10,000  (A9044-2 mL) | Millipore Sigma  Burlington, MA |
| NeuN  (66836-1-Ig) | Mouse monoclonal  1:500 | Proteintech, Rosemont, IL | Rabbit anti-mouse IgG HRP 1:10,000  (A9044-2 mL) | Millipore Sigma  Burlington, MA |
| GFAP  (60190-1-Ig) | Mouse monoclonal  1:1000 | Proteintech, Rosemont, IL | Rabbit anti-mouse IgG HRP 1:10,000  (A9044-2 mL) | Millipore Sigma  Burlington, MA |
| IBA1  (66827-1-Ig) | Mouse monoclonal 1:1000 | Proteintech, Rosemont, IL | Rabbit anti-mouse IgG HRP 1:10,000  (A9044-2 mL) | Millipore Sigma  Burlington, MA |
| APP (6E10)  (NBP2-62566) | Rabbit monoclonal  1:1000 | Novus Biologicals, Minneapolis, MN | Goat anti-rabbit IgG HRP 1:10,000  (A9169-2 mL) | Millipore Sigma  Burlington, MA |
| Tau  (13-6400) | Mouse monoclonal 1:1000 | Thermo Fisher Scientific, MA | Rabbit anti-mouse IgG HRP 1:10,000  (A9044-2 mL) | Millipore Sigma  Burlington, MA |
| GAPDH  (2118) | Rabbit monoclonal  1:3000 | Cell Signaling, MA | Goat anti-rabbit IgG HRP 1:10,000  (A9169-2 mL) | Millipore Sigma  Burlington, MA |
