## Supplementary material for "MiRNA-501-3p and MiRNA-502-3p: A Promising Biomarker Panel for Alzheimer’s Disease": SI Table 7

**Supplementary Table 7. Oligonucleotide sequences of primers used for quantitative reverse transcription-polymerase chain reaction analysis**

| **Gene(s)** | **Sequence(s)** |
| --- | --- |
| Hsa-APP | F- 5^'^ GCCGATGATGACGAGAGAGG 3^'^ |
|  | R- 5^'^ GGGTACTGGCTGCTGTTGTA 3^'^ |
| Hsa-Tau | F- 5^'^ AAAGCCAAGACAGACCACGG 3^'^ |
|  | R- 5^'^ AGCTTCTGCAGGTCGACTCAC 3^'^ |
| GAPDH | F- 5^'^ GCACCGTCAAGGCTGAGAAC 3^'^ |
|  | R- 5^'^ TGGTGAAGACGCCAGTGG 3^'^ |
| U6 SnRNA | F- 5^'^ CGCTTCGGCAGCACATATACTAA 3^'^ |
|  | R- 5^'^ TATGGAACGCTTCACGAATTTGC 3^'^ |
| miR-502-3p | F- 5^'^ AATGCACCTGGGCAAGGATTCA 3^'^ |
| miR-501-3p | F- 5^'^ AATGCACCCGGGCAAGGATTCT 3^'^ |
