## Supplementary material for "MiRNA-501-3p and MiRNA-502-3p: A Promising Biomarker Panel for Alzheimer’s Disease": SI file

#### Vector Summary

| | |
|---|---|
| **Vector ID** | VB010000-9829sne |
| **Vector Name** | pRP[Exp]-CAG>EGFP |
| **Vector Size** | 4801 bp |
| **Vector Type** | Mammalian Gene Expression Vector |
| **Inserted Promoter** | CAG |
| **Inserted ORF** | EGFP |
| **Plasmid Copy Number** | High |
| **Antibiotic Resistance** | Ampicillin |
| **Cloning Host** | Stbl3 (or alternative strain) |

#### Vector Map

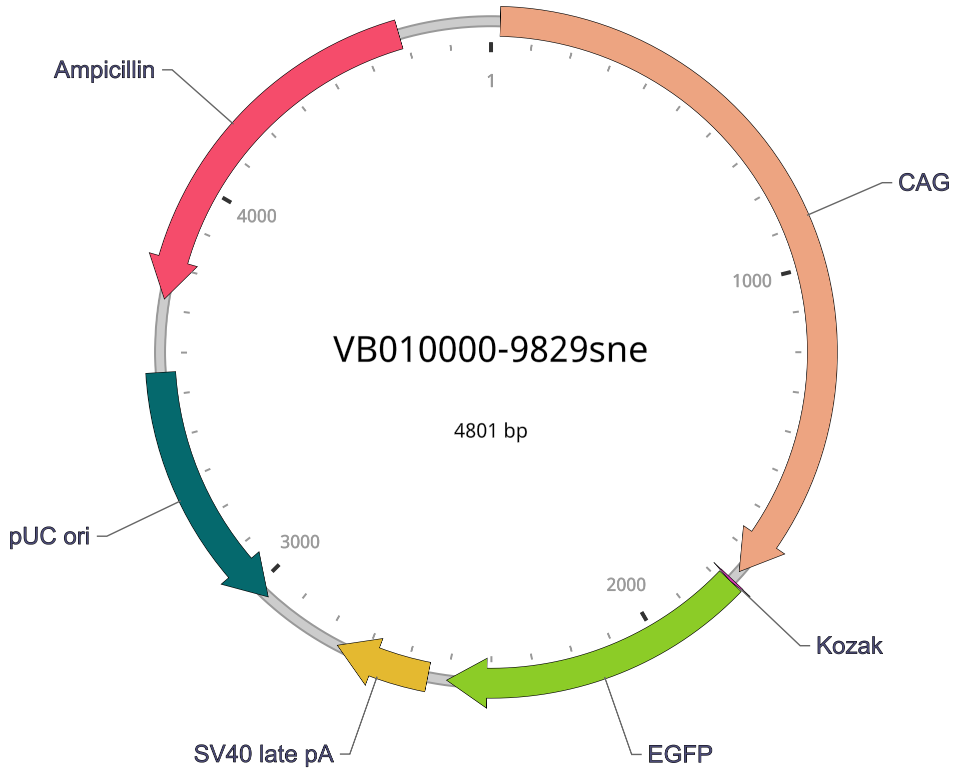

addgene

### Sequence Analyzer: pCAX APP Swe/Ind Sequencing Result

**Download:** GenBank File | SnapGene File | File Help

Map Sequence Enzymes Features Primers BLAST Analyzer Guide

#### Map View

Instructions

Created with SnapGene®

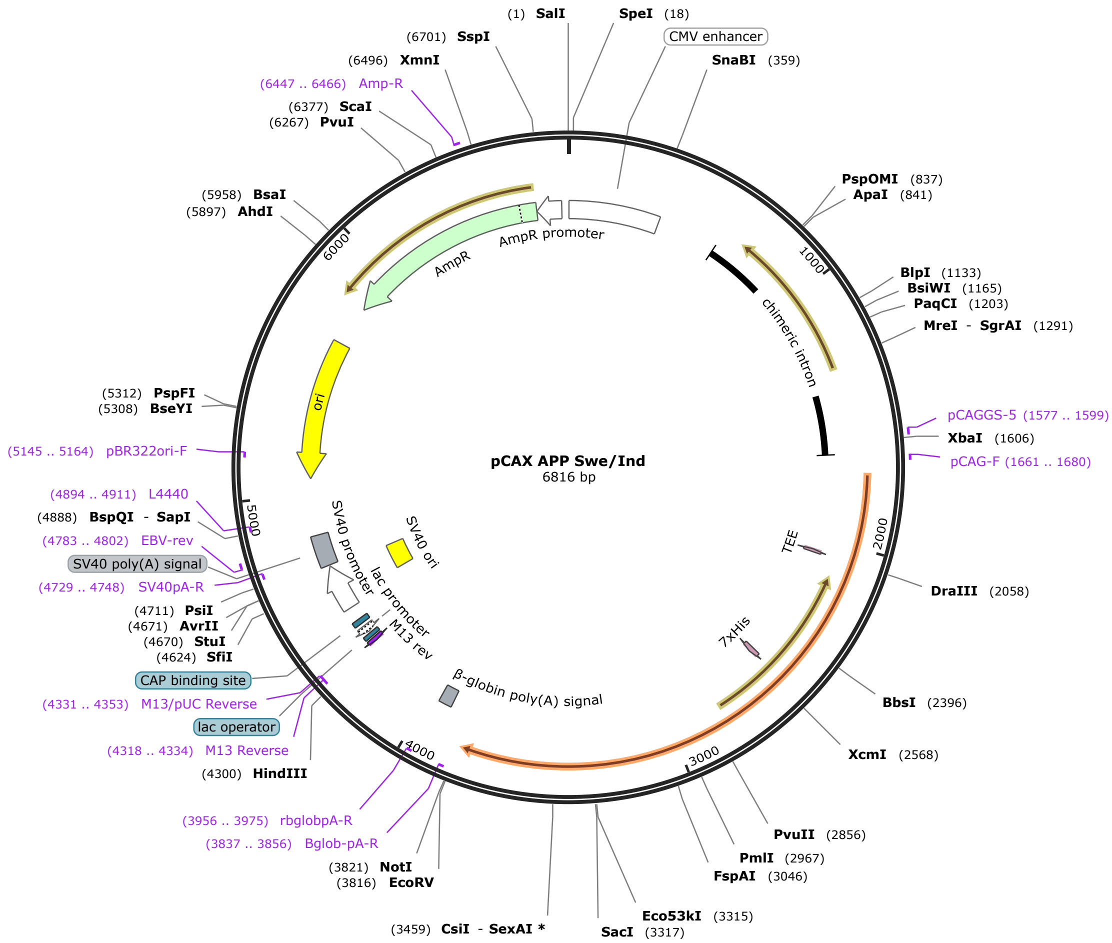

addgene

### Sequence Analyzer: pAAV-FLEX-P301L Tau Sequencing Result

**Download:** GenBank File | SnapGene File | File Help

Map Sequence Enzymes Features Primers BLAST Analyzer Guide

#### Map View

Instructions

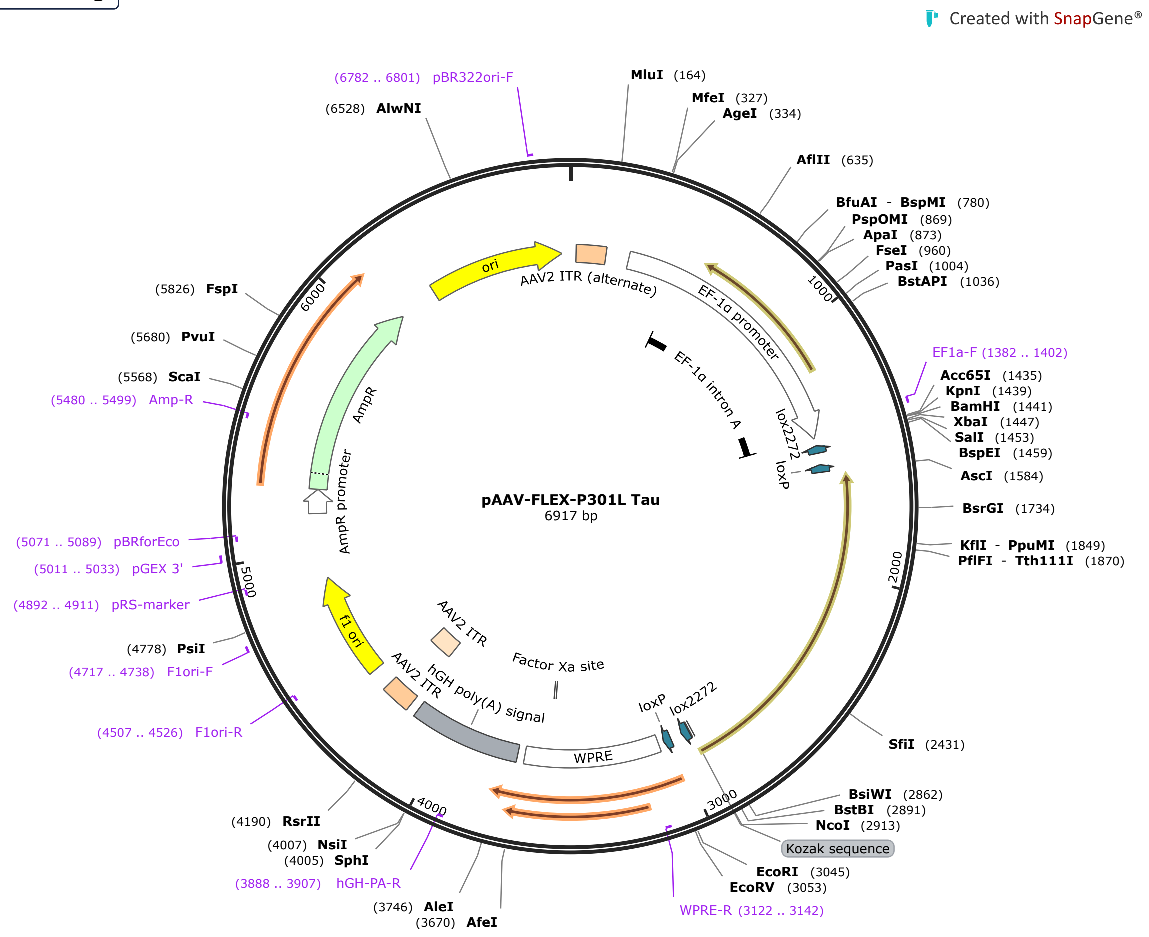
